# Cellular heterogeneity in the developing forebrain masks transcriptional outcomes and principles of *Evf2* enhancer lncRNA-*Dlx5/6UCE*-gene guidance

**DOI:** 10.1101/2024.02.20.581236

**Authors:** Edward Li, Abhijit Chakraborty, Sara J. Kohtz, Ivelisse Cajigas, Fion Shiau, Robert J. Vassar, Brian S. Clark, Jhumku D. Kohtz

## Abstract

During mouse embryonic brain development, the *Evf2* ultraconserved enhancer (UCE) lncRNA guides the Dlx5/6UCE to ∼129 sites across chr6. However, previous work identified only 4 transcriptionally regulated targets associated with *Evf2*-Dlx5/6UCE enhancer-gene guided sites (EGGs), raising questions about the significance of the majority of *Evf2*-EGGs. Here, single cell transcriptomics (scRNAseq) shows that *Evf2*-EGGs on chr6 coincide with subpopulation-specific *Evf2* transcriptional targets, revealing far greater alignment between EGGs and transcriptional targets than previously reported. Surprisingly, subpopulation-specific *Evf2* regulated gene networks in embryonic progenitors predict adult synaptic and seizure defects. *Evf2* regulation of EGGs to gene bodies (GB, Dlx5/6UCE locations within ±5kb of the target gene) divides chr6 into short-range (<10Mb distant), highly activated genes, and long/super-long-range (10-129Mb), moderately repressed genes. Clustering of *Evf2* transcriptionally regulated chr6 targets in populations where *Evf2* is first activated supports that *Evf2*-EGG transcriptional effects can occur from EGG shifts as far as 3Mb, uncoupling enhancer shift distance and transcriptional direction from transcriptional outcome. *Evf2* RNA binding sites (RBSs) divide chr6 into 4 major regions, consistent with a role for RBS chromosomal spacing in long-and super-long-range EGG selectivity. Surprisingly, 95% of 147 RBSs genome-wide potentially form inter-chromosomal DNA loops. *Evf2*-regulated combinatorial recruitment of *Evf2*-ribonucleoproteins at EGGs and RBSs, together with effects on homeodomain transcription factor DNA motif recognition, support a novel model of lncRNA directed multi-modal EGG selectivity during intra- and inter-chromosomal gene regulation.

## Introduction

With the discovery that DNA regulatory sequences (enhancers) selectively regulate genes across megabase distances of the genome, it is now clear that co-regulation does not require co-linear gene organization, but rather 3D organization. Understanding how enhancers select specific target genes, while skipping others has become a major focus in the field. Multiple 3D gene organizational mechanisms have been proposed, including formation of intra-chromosomal and inter-chromosomal enhancer-gene 3D hubs, loop extrusion through CTCF and cohesin recruitment, R-loop formation and CTCF-RNA binding (Lim and Levine, 2021; Uyehara and Apostolou, 2023).

Initial studies on short enhancer non-coding RNAs (eRNAs) demonstrated enhancer regulating activities (Orom et al., 2010; Orom and Shiekhattar, 2011) and short-range chromosome looping activity (Lai et al., 2013). However, genome-wide roles for enhancer long non-coding RNAs (e-lncRNAs) as organizers have also been proposed (Mattick, 2023; Mattick et al., 2023), supported by enrichment at chromosomal loop anchors, increased frequency of interactions with promoters, RNA Pol II and YY1 binding, and G-quadruplexes (Hou et al., 2019). A human atlas of 45,411 eRNAs (HeRA) from 9577 samples across 54 human tissues (Zhang et al., 2021), and a compendium of eRNA functional assays highlights their potential significance (Yao et al., 2022) and dual regulatory functions at the level of RNA and DNA.

Functional roles for individual lncRNAs and enhancer RNAs (eRNAs) support roles in 3D organization. Investigations into lncRNA and e-lncRNA regulators of 3D organization reveal both intra-[Xist (Nora et al., 2012), *ThymoD* lncRNA (Isoda et al., 2017), *Evf2* (Cajigas et al., 2018)] and inter-[Firre (Hacisuleyman et al., 2014), *DRReRNA* (Tsai et al., 2018)] chromosomal mechanisms. The *Xist* lncRNA is among the most well characterized lncRNAs that regulates intra-chromosomal 3D organization (Giorgetti et al., 2016; Nora *et al*., 2012). Proteomic analysis of the *Xist*-RNP has revealed roles of several RNPs (Chen et al., 2016; Chu et al., 2015; Dossin et al., 2020; Minajigi et al., 2015; Yi et al., 2020). Our work on *Evf2* identified nuclear e-lncRNA cloud formation with Dlx homeodomain transcription factors, and trans-regulation of enhancer activity through ultraconserved sequences (Feng et al., 2006). *Evf2* and *Xist* exhibit several similarities, both forming RNA clouds at sites of transcription initiation, altering chromosome topology and repressing genes through effects on chromatin remodeling (Cajigas *et al*., 2018; Cajigas et al., 2015; Feng *et al*., 2006; Giorgetti *et al*., 2016; Jegu et al., 2019; Nora *et al*., 2012). The Xist-RNP^85^ contains 16 proteins (Chu *et al*., 2015), shared with the *Evf2-* RNP^87^ (Cajigas *et al*., 2015), including Smarca4, the catalytic subunit of BAF chromatin-remodeling complex, and 3D organizer cohesin (Smc1a and Smc3) (Cajigas *et al*., 2015; Minajigi *et al*., 2015). *Evf2* RNA- and G-rich RNA-Smarca4 interactions inhibit chromatin remodeling activity through inhibition of ATPase activity (Cajigas *et al*., 2015), a mechanism also shared during *X-*inactivation (Jegu *et al*., 2019). Furthermore, *Evf2*-dependent recruitment of cohesin near key topologically controlled sites supports involvement in *Evf2-Dlx5/6UCE* long-range gene guidance (Cajigas *et al*., 2018); both intra-(*ThymoD* lncRNA (Isoda *et al*., 2017)) and inter-(*DRReRNA* (Tsai *et al*., 2018)) chromosomal RNA regulators also recruit cohesin, supporting shared lncRNA-3D mechanisms.

While our previous work showed that *Evf2* guides *Dlx5/6UCE* to sites across an entire chromosome, in part, through Sox2 and cohesin recruitment (Cajigas et al., 2021; Cajigas *et al*., 2018), the significance of the majority of Dlx5/6UCE guidance events was not known. In addition, despite the demonstration of *Evf2* RNA cloud formation at transcriptionally regulated target genes, directly bound Evf2 sites, distinguishing between direct and cascade mechanisms were not reported. In this work, we combine scRNAseq, ChIRPseq, and Cut&Run with previously reported Dlx5/6UCE chromosome conformation capture (4Cseq) and Cut &Run/ChIPseq datasets (Cajigas *et al*., 2021; Cajigas *et al*., 2018), to identify a role for 1D and 3D enhancer-gene relationships in *Evf2* e-lncRNA-mediated selectivity in transcriptional regulation. Taken together, these data link multiple *Evf2* regulatory events, including *Evf2* RNA-DNA binding, RNP recruitment, transcription factor-DNA motif recognition, and Dlx5/6UCE guidance to selectively regulate gene transcription.

## Results

### Cellular heterogeneity masks the extent of *Evf2* transcriptional regulation in mouse E13.5 ganglionic eminences

Chromosome conformation capture (4Cseq) studies using Dlx5/6UCE as the bait identified 129 *Evf2*-regulated EGGs (located within 50kb of gene targets on chr6) (Cajigas *et al*., 2018). However, bulk analysis (microarray) of E13.5GEs identified only 6 *Evf2* regulated chr6 genes, 2 adjacent genes (Dlx5 and Dlx6), and 4 long-range EGG-associated genes (Cajigas *et al*., 2018). One possibility was that EGG regulation poises genes for regulation later in development (Ghavi-Helm et al., 2014). However, heterogeneity of *Evf2* RNA nuclear cloud numbers and sizes (Berghoff et al., 2013; Cajigas *et al*., 2021), *Evf2*-regulated Sox2 protein pool/condensate heterogeneity (Cajigas *et al*., 2021), *Evf2*-regulated enhancer-gene distance heterogeneity (Cajigas *et al*., 2018), and transcriptional heterogeneity among E13.5GE subpopulations in mice and primates (Mayer et al., 2018; Schmitz et al., 2022), supported the possibility that E13.5GE cellular heterogeneity masks the extent of *Evf2* transcriptional effects. In order to determine the full extent of *Evf2* transcriptional regulation, we used 10X genomics scRNAseq (Fig 1A-F, Fig S1) to compare gene expression in single E13.5GE cells from wildtype (*Evf2^+/+^*) and mice with a targeted transcription stop in Evf-exon 1, preventing *Evf2* expression, (*Evf2^TS/TS^*) (Bond et al., 2009)). Cell type annotations are determined using marker specificity within clusters of cells, as determined by genesorteR (https://github.com/mahmoudibrahim/genesorteR). Subpopulations corresponding to previously annotated cell types (Mayer *et al*., 2018), and cycling cells (ventricular zone [VZ], subventricular zone [SVZ]) are shown in UMAPs for each genotype (Fig 1A, B). *Evf2* is activated and co-expressed with Dlx5/6 in cells transitioning from VZ to SVZ (Fig 1C, D), consistent with previous RNA in situ analysis (Berghoff *et al*., 2013; Bond *et al*., 2009; Feng *et al*., 2006). The left branch (L) of the UMAP identifies *Evf2*(+) populations in cycling progenitors (SVZ) and differentiated neuronal populations (Lhx6, Lhx6/8) (Fig 1A), whereas the right branch (R) identifies *Evf2*(-) (NeuroG2, NeuroD6, and Lhx1/5) populations that have exited the SVZ. In the left branch, Isl1+ cells express low levels of *Evf2* and high levels of Dlx6, consistent with *Evf2* (also known as Dlx6OS1) - mediated Dlx6 anti-sense gene repression (Cajigas *et al*., 2018).

**Figure 1.**
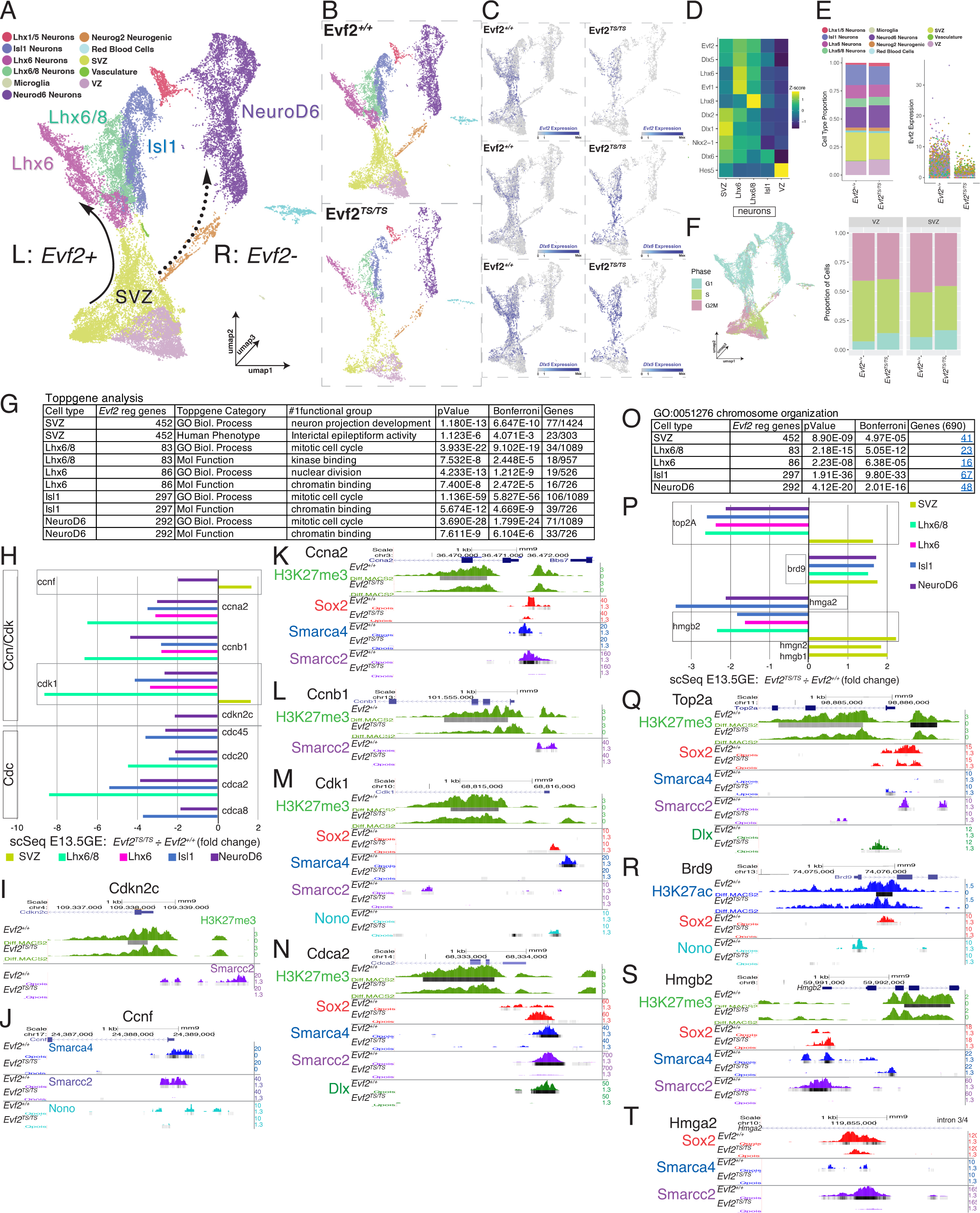
Single cell sequencing of E13.5 ganglionic eminences reveals *Evf2* regulated shared and unique functional gene networks among subpopulations, with the subventricular zone (SVZ) predicting adult effects on epileptiform activity genes. 10Xgenomics scRNAseq datasets were generated from sequencing *Evf2^+/+^* and *Evf2^TS/TS^* E13.5GEs. **A.** UMAP of gene expression of E13.5GEs at single cell resolution identifies subpopulations (different colors) and two major branches relevant to *Evf2* regulation (left, L (*Evf2*+), and right, R (*Evf2*-)). *Evf2* is activated in cells transiting from cycling progenitor zones (ventricular zone, VZ, light pink) and subventricular zone (SVZ, yellow). Arrows represent transitions from the SVZ to differentiated populations: L: *Evf2*+ (Lhx6, Lhx6/8, Islet1) and R:*Evf2*-(NeuroD6) populations. **B-C.** UMAPs comparing *Evf2^+/+^* and *Evf2^TS/TS^* subpopulations. *Evf2* is co-expressed with Dlx5 and Dlx6 in the left branch. **D**. Heatmap confirming the relative expression of genes that define specific E13.5GE subpopulations, with *Evf2*/*Evf1*/Dlx family (Dlx1,2,5,6). **E.** left: Relative proportions of subpopulations compared between *Evf2^+/+^* vs *Evf2^TS/TS^* are unaltered, right: *Evf2* expression at the single cell level is plotted. **F.** (Left): UMAP of cell cycle phases (G1, S, G2/M); (Right) *Evf2^+/+^* vs *Evf2^TS/TS^* proportions of cells in specific cell cycle phases. **G**. Toppgene analysis of SVZ and SVZ-derived subpopulations, identifying shared and distinct functional groups. **H**. In *Evf2^TS/TS^*, cell cycle target genes (9) decrease in differentiated populations, while 2/9 increase in the subventricular zone. **I-N.** *Evf2* regulation of histone modification (H3K27me3), and/or *Evf2*-RNP recruitment (Smarca4, Smarcc2, Sox2, Dlx, Nono) at cell cycle genes, as indicated in peaks comparing *Evf2^+/+^* vs *Evf2^TS/TS^*. **O**. Toppgene analysis of SVZ-derived subpopulations identifying genes involved in chromosomal organization. **P.** In *Evf2^TS/TS^*, genes involved in chromosomal organization are differentially regulated among subpopulations. **Q.-T** *Evf2* regulation of histone modification (H3K27me3, H3K27ac), and/or *Evf2*-RNP recruitment (Smarca4, Smarcc2, Sox2, Dlx, Nono) at genes involved in chromosome organization, as indicated in peaks comparing *Evf2^+/+^*vs *_Evf2TS/TS_*.

Differential scRNAseq comparing *Evf2^+/+^* vs. *Evf2^TS/TS^*E13.5GE subpopulations reveal a far greater extent of *Evf2* transcriptionally regulated target genes than previously identified (Fig 1). In *Evf2*(+) UMAP left branch subpopulations, there are 559 SVZ [293(activated)/265(repressed)], 107 Lhx6, and 101 Lhx6/8 *Evf2* transcriptional targets (cutoffs of >100cells, >1.5 fold change, qval<10^-5^). *Evf2* transcriptional targets are detected in populations that either no longer express *Evf2*, or express *Evf2* at low levels after exiting the SVZ (362 (Isl1), 364 (NeuroD6), and 60 (NeuroG2)), supporting either a requirement of *Evf2* in SVZ progenitors prior to differentiation, and/or cell non-autonomous effects. Despite extensive gene expression effects within subpopulations, *Evf2* does not appear to affect overall cell type or cell cycle proportions in SVZ (Fig 1E, F) arguing against a role in fate specification or cell cycle control. Toppgene analysis of *Evf2* regulated gene regulatory networks (GRNs) identifies both common and unique GO functional networks (Fig 1G). The SVZ is unique in that the number 1 functional *Evf2* SVZ GRNs are involved in neuronal development (GO Biological Process) and epilepsy (Human phenotype), consistent with developmentally regulated synaptic defects and increased seizure susceptibility that appear in adult mice lacking *Evf2* (Bond *et al*., 2009; Cajigas *et al*., 2018). In both L and R differentiated neuronal subpopulations (Lhx6, Lhx6/8, Isl1, NeuroD), mitotic cell cycle genes or nuclear division are the number 1 *Evf2* regulated GO Biological Process. In *Evf2^TS/TS^* differentiated populations, but not SVZ, cell cycle regulators are decreased, supporting effects on transitory cells (cells expressing both cell cycle and differentiated genes) (Fig 1H). Previously reported native ChIPseq, crosslinked ChIPseq, Cut&Run (Cajigas *et al*., 2018) combined with newly generated Cut&Run profiles (*Evf2* RNPs Smarcc2 and Nono) show that *Evf2* regulates H3K27me3 and/or *Evf2* RNP recruitment to cell cycle genes, (Fig 1I-J). Although bulk E13.5 GE tissues are used for ChIPseq/Cut&Run methods, *Evf2*-regulated RNP recruitment changes are detected at targets regulated in a single subpopulation (NeuroD6/Cdkn2c, Fig 1I), as well as at targets with subpopulation-specific effects (Ccnf/Cdk1, increased in SVZ, decreased in differentiated populations, Fig 1J, M). Alignment of *Evf2* regulated RNP binding and histone modifications at scRNAseq gene targets supports transcriptional rather than post-transcriptional mechanisms.

The number 1 GO molecular function (chromatin binding) is shared among *Evf2* regulated Lhx6/Isl1/NeuroD6 GRNs, while the GO Biological process, chromosome organization (Fig 1O-T, Fig S1D, Venn diagram) is present in the 5 major SVZ-derived subpopulations, supporting transcriptional regulation of chromosome organizers during *Evf2*-regulated chromosome organization (Cajigas *et al*., 2021; Cajigas *et al*., 2018). Overall, transcripts encoding chromosome organizers increase in *Evf2^TS/TS^* SVZ, and decrease in differentiated subpopulations, except for Brd9, which increases independent of differentiation (Fig 1O). *Evf2*-regulates H3K27me3/H3K27ac and *Evf2*-RNP binding at genes involved in chromosome organization, supporting transcriptional rather than post-transcriptional effects. Thus, subpopulation-specific *Evf2* transcriptional regulation of key chromosome organizers supports the idea that heterogeneity of chromosomal organization in E13.5GEs is established, in part, through transcriptional regulation during neuronal differentiation.

### Transcriptional gene targets align with multimodal *Evf2* EGGs and RNP binding signatures

We next asked how *Evf2* transcriptionally regulated targets align with *Evf2* regulated EGGs on mouse chr6, combining differential scRNAseq results in E13.5GE subpopulations and *Dlx5/6UCE-4Cseq* results that identified 129 *Evf2* regulated EGGs (Cajigas *et al*., 2021; Cajigas *et al*., 2018). We first focused on Dlx5/6UCEs located within 5kb of gene bodies (GB), as the closest *Evf2* regulated EGG-GB shifts are expected to have the highest probability of transcriptional effects. EGG-GB shifts align with 16 transcriptionally regulated targets on chr6 across 5 subpopulations (colored boxes, Fig 2A); 9/16 are also regulated by Sox2 (bi-colored boxes, previously identified by *Dlx5/6UCE-4Cseq* (Cajigas *et al*., 2021)). Grey boxes (Fig 2A) identify 3 transcriptionally regulated targets associated with *Evf2* independent-*Dlx5/6UCE* interactions. These 19 *Evf2* transcriptionally regulated genes associated with *Dlx5/6UCE* locations within 5kb of the GB are referred to as *Dlx5/6UCE*-GB^19^. Analysis of *Dlx5/6UCE*-GB^19^ targets reveals differences in transcriptional effects based on relative positions on *chr6* (fixed distances from *Dlx5/6UCE* in one dimension (1D)). Specifically, *Evf2* activates short-range genes located within 10Mb of *Dlx5/6UCE* (Ccdc132-FoxP2) by ∼8.8 fold, and represses long- and super-long-range targets located >10Mb from *Dlx5/6UCE* (Gm15594-Grin2b) by ∼1.8fold, dividing chr6 into *Evf2*-regulated activated and repressed domains. Importantly, although 4Cseq is performed on bulk E13.5GE chromatin, *Evf2*-regulated EGGs associated with transcriptionally regulated subpopulation-specific scRNAseq GRNs are identified.

**Figure 2.**
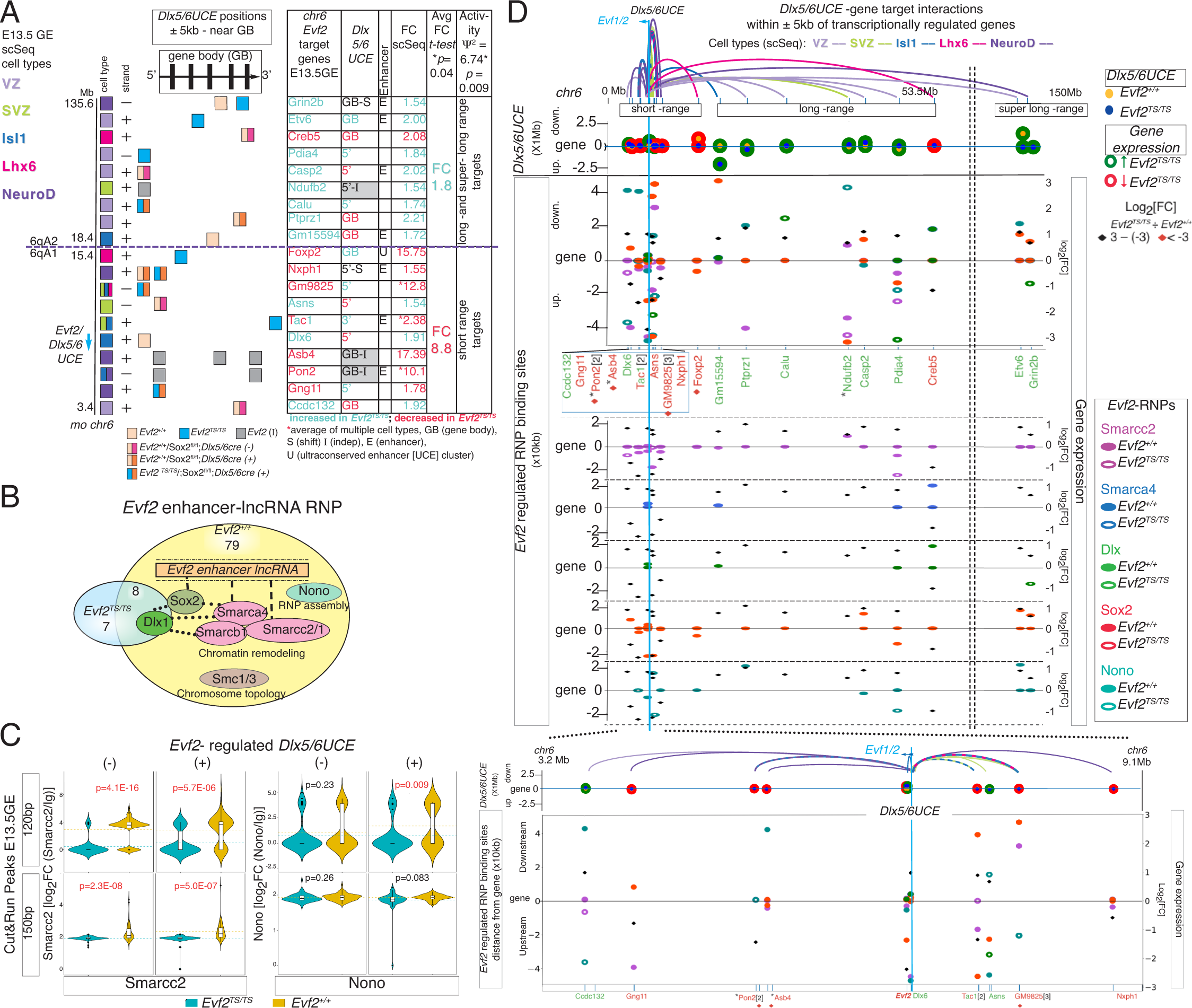
*Evf2* guidance of Dlx5/6UCE to gene bodies (±5kb) divides chr6 into transcriptionally activated (short-range) and repressed (long-super long-range) domains. Differential scRNAseq analysis of *Evf2* regulated genes located on chr6 (*Evf2^TS/TS^÷ Evf2^+/+^)* identifies cell type specificity, represented according to color (VZ, SVZ, Isl1, Lhx6 and Neurod6). **A**. *Evf2* target gene locations on chr6 from 3.5Mb (Ccdc132) -135Mb (Grin2b), vertically represented with strand information, colored boxes corresponding to cell types. *Evf2* transcriptionally regulates 19 genes on chr 6 where Dlx5/6UCE is located within ±5kb of the gene body (GB). *Evf2* guides Dlx5/6UCE to 16/19 GBs. 3D-Dlx5/6UCE positional shifts relative to GBs in the presence (*Evf2^+/+^*, pink boxes) and absence (*Evf2^TS/TS^*, blue boxes) of *Evf2*, co-regulated by Sox2 (bi-colored boxes), or grey boxes (independent of *Evf2*). Genes associated with enhancers, E, or an ultraconserved sequence, U (Foxp2) are identified. Chr6 qA1/A2 (dashed line) divides chr6 into genes that decrease (red) and genes that increase (blue) in *Evf2^TS/TS^*, average fold differences, two tailed *t*-test (p=0.04), regional separation of increased/decreased targets, (ψ=6.74, p=0.009). **B.** The *Evf2* RNP^87^ (yellow circle) consists of diverse RNA binding proteins (dashed lines) and protein-protein interactions (dotted lines), identified Smarcc2 and Nono as *Evf2* RNPs, based on (Cajigas *et al*., 2015). **C.** Violin Plots compare *Evf2* regulated RNP binding in E13.5GEs (*Evf2^TS/TS^* vs. *Evf2^+/+^*). Cut&Run 120bp fragments (directly bound) and 150bp fragments (indirectly bound) profiles of Smarcc2 and Nono peaks at *Evf2* positively (+) and negatively (-) regulated *Dlx5/6UCEins* (4Cseq) across *chr6*. Two tailed Student’s *t*-test, p<0.05 in red, blue (*Evf2^TS/TS^*) yellow (*Evf2^+/+^*). **D.** *Evf2*-EGG-GB plots align *Dlx5/6UCE* shifts at each *chr6* transcriptionally regulated gene target and distances of associated *Evf2* regulated *Evf2-*RNP binding sites. Colored loops highlight *Evf2* differentially regulated scRNAseq gene targets on chr6 (*Evf2^TS/TS^÷Evf2^+/+^)* in E13.5GE subpopulations (VZ, SVZ, Isl1, Lhx6 and Neurod6) coincident with *Evf2*-Dlx5/6UCE guidance: Top X-axis, chr6: 0-150Mb, panel focus below: Zommed X-axis, chr6: 3.2-9.1Mb. Dlx5/6UCE Y-axis (x1Mb up (upstream) and down (downstream) distances from gene (0)): *Evf2(+)*regulated *Dlx5/6UCE* position (yellow dots), *Evf2(-)*regulated Dlx5/6UCE position (blue dots). Transcriptional outcomes of *Evf2* loss (*Evf2^TS/TS^*): shifts associated with decreased gene expression (red rings), shifts associated with increased gene expression (green rings). Right Y axis (gene expression): log_2_ (fold change in gene expression), black diamonds (-3 - +3), red diamonds (<-3). Y-axis *Evf2*-regulated RNP binding sites (x10kb distances from gene (0)) (Smarcc2, Smarca4, Dlx, Sox2, Nono, Smc1): solid ovals (*Evf2(+)* regulated binding), rings (*Evf2(-)* regulated binding). Blue line indicates location of Dlx5/6UCE/*Evf1/2* on chr6.

In order to further understand the mechanism of *Evf2*-EGG regulation, we investigated the relationship between *Evf2*-EGGs and the *Evf2* RNP. The *Evf2* RNA cloud is a scaffold for the assembly of a diverse Dlx-bound, ribonucleoprotein complex initially identified by proteomic analysis of E13.5GEs (*Evf2*-RNP^87^) (Fig 2B); *Evf2* RNA directly binds chromatin remodelers Smarca4, Smarcc2/1, and binds the Sox2 homeodomain TF through promiscuous RNA-protein interactions, while protein-protein interactions indirectly link *Evf2* RNA to the Dlx homeodomain TF (Cajigas *et al*., 2021; Cajigas *et al*., 2018; Cajigas *et al*., 2015). Previous work showed that *Evf2* recruitment of diverse *Evf2*-RNPs (Sox2, Dlx, Smarac4, Smc1a, Smc3) are enriched at *Dlx5/6UCE* regulated sites, with the *Evf2-*Sox2 RNP contributing to EGG selectivity on chr6 (Cajigas *et al*., 2021). We expanding our previous analysis of *Evf2*-RNP recruitment by performing Cut&Run of Smarcc2 (a chromatin remodeler with promiscuous RNA binding properties similar to Smarca4, (Cajigas *et al*., 2015)), and Nono (the Neat-1 arcRNA binding, paraspeckle organizer (Yamazaki et al., 2018)). Violin plots identify significant differences in the numbers of Smarcc2 bound peaks at Dlx5/6UCE (+) and (-) regulated sites in the presence of *Evf2* (*Evf2^+/+^*, yellow) compared to absence (*Evf2^TS/TS^*, blue) (Fig 2C). While *Evf2* also regulates Nono binding at Dlx5/6UCE (+) regulated sites, regulation is limited compared to Smarcc2, identified in directly bound sites (120bp fragments), thereby predicting differential roles for Smarcc2 and Nono.

We next created EGG plots in order to visualize *Dlx5/6UCE*-GB^19^ targets on chr6 (1D fixed positions, X-axis) in relation to *Evf2*-regulated *Dlx5/6UCE* shifts (3D shifted positions, Y-axis, (Fig 2D) and *Evf2*-regulated RNP (Smarcc2, Smarca4, Dlx (1/2/5/6), Sox2, Nono) binding. Yellow and blue solid circles indicate *Dlx5/6UCE* positions relative to GBs in 3D (*Evf2^+/+^(*yellow solid circle) and *Evf2^TS/TS^* (blue solid circle)), while green (increased transcription) and red (decreased transcription) rings indicate transcriptional outcomes in *Evf2^TS/TS^* (Fig 2D). EGG plots identify at least one or more *Evf2*-regulated RNPs within ±50kb of GBs (Fig 2D [chr6 0-150mb], inset and Fig S2: chr6; 3-9Mbregion [Ccdc132-Nxph1]), data consistent with diverse *Evf2* RNP roles in chromosome organization, chromatin remodeling and transcriptional regulation (Cajigas *et al*., 2021; Cajigas *et al*., 2018; Cajigas *et al*., 2015).

Since *Evf2* is activated in cells exiting from the VZ to SVZ (Bond *et al*., 2009; Feng *et al*., 2006), and expression levels are highest in SVZ progenitors, SVZ transcriptional targets constitute the most likely direct targets of *Evf2* regulation. However, transcriptional effects of only 4 SVZ EGG-GB targets are detected, raising the possibility that long-distance EGG shifts may also contribute to gene regulation in the SVZ. We next compared EGG plots of chr6 *Evf2* regulated target genes in SVZ (29 genes, Fig 3A) with Dlx+ populations (UMAP left branch differentiated subpopulations derived from the SVZ, 23 genes, Fig 3B), aligning genes on chr6 (1D fixed positions, X-axis) and their nearest *Evf2* regulated Dlx5/6UCE shifts (3D dynamic positions, Y-axis). In the SVZ, as target gene distances from Dlx5/6UCE increase in 1D, EGG-3D distances also increase, averaging 380kb for genes located between Tac1-Lsm5 and 5.1Mb for genes located between Med2l1-Fam60a, a 13.5-fold increase (3A). Arrows indicate the direction of each EGG shift (in the absence of *Evf2*) with respect to the closest transcriptionally regulated target gene (GBs). In the SVZ, *Evf2* regulated long-distance EGG shifts in relation to GBs range between 5kb-15.4Mb, characterized by various positional changes (upstream, downstream, and across-GB). The identification of gene clusters (≥2 genes) associated with Dlx5/6UCE shifts that are equidistant (two-color dashed arrows), or similar (to within ±50kb, solid arrows), support co-regulatory 3D-causing transcriptional changes. The largest group of equidistant genes consists of the furthest 4-gene cluster (Eno-Cdca3-Rad51ap1-Prmt8), in which *Evf2* shifts Dlx5/6UCE towards the gene targets, increasing Eno and Prmt8, and decreasing Cdca3 and Rad51ap1 expression. Therefore, even though *Evf2* shifts Dlx5/6 closer to a gene, transcriptional effects can be positive or negative, a characteristic also observed for short distance EGG-GB shifts. These results support the idea that *Evf2* regulated Dlx5/6UCE-3D shift distances and direction can be decoupled from transcriptional outcome.

**Figure 3.**
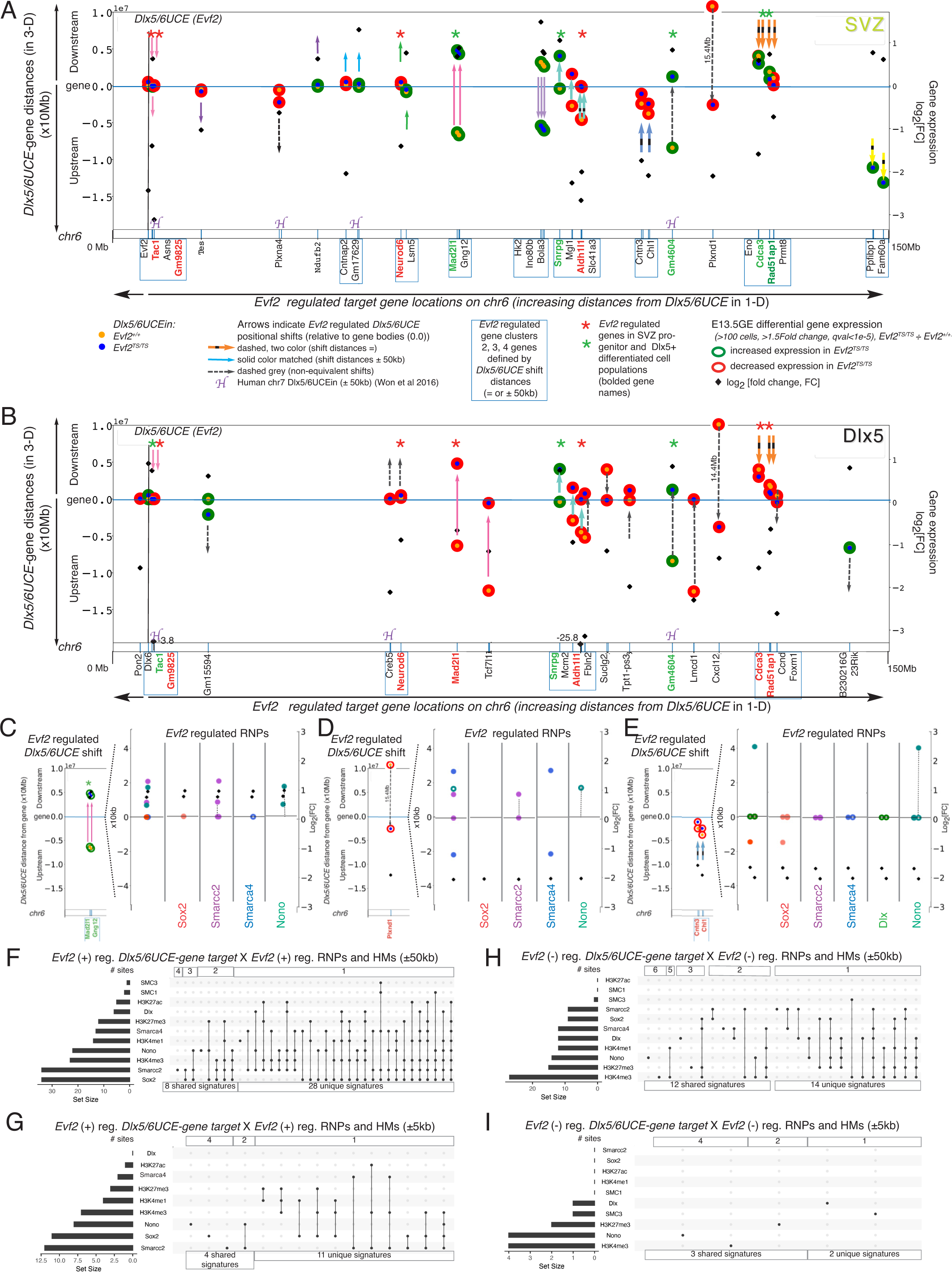
Multimodal *Evf2* Dlx5/6UCE guidance in clusters of genes coincident with RNP binding/histone modification effects and transcriptional regulation during differentiation. **A-B**. *Evf2*-EGG plots of left branch (A) SVZ progenitors and (B) Dlx5+ differentiated (Lhx6, Lhx6/8, Islet1) cell types at transcriptionally regulated target genes located on chr6 (X-axis, 0-150Mb: gene locations), Dlx5/6UCE 3D-distances from gene bodies (0) on left Y axis (X10Mb), gene expression (log2[FC]) on right Y axis. Arrows indicate *Evf2*-guided Dlx5/6UCE shift directions towards or away from gene bodies; arrow colors reflect shared shift distances, (as indicated in legend). Gene clusters (2-4) are identified by equidistant Dlx5/6UCE shifts (dashed two color arrows) or shifts within 50kb (solid color arrows) are boxed. Conservation with human developing forebrain chr7 Dlx5/6UCE interaction sites identified by HiC (Won *et al*., 2016) are labelled as H. *Evf2(+)*regulated *Dlx5/6UCE* position (yellow dots), *Evf2(-)*regulated Dlx5/6UCE position (blue dots), distance from the gene (0). Transcriptional outcomes in *Evf2^TS/TS^*: shifts associated with decreased gene expression (red rings), shifts associated with increased gene expression (green rings). Right Y axis: log_2_ (fold change in gene expression), black diamonds (-3 - +3), and two off-scale targets in Dlx+ are marked (Gm9825, -3.8, Aldh1l1, -25.8). Stars indicate 9 genes that are regulated in both SVZ and Dlx5+ cell types (green, increased, red, decreased). **C-E**. Examples of multimodal *Evf2*-EGGs (located ±10Mb from gene target) coincident with *Evf2* regulated multi-RNP binding of Sox2, Smarcc2, Smarca4, Nono (±50kb) and changes in transcriptional outcomes in the SVZ. **C.** Similar distance (pink arrows) *Evf2*-Dlx5/6UCE guidance across Mad2l1 and Gng12, with Sox2/Smarcc2/Smarca4 RNP recruitment effects at GBs, and asymmetric Smarcc2/Nono downstream effects. **D.** 15.4 Mb (grey dashed arrow) *Evf2*-Dlx5/6UCE guidance across Plxnd1, with RNP effects: GB (Smarcc2), upstream and downstream (Smarca4), asymmetric downstream (Smarcc2/Nono). **E.** Upstream *Evf2*-Dlx5/6UCE guidance, with RNP effects: GB (Sox2, Smarcc2/Smarca4, Dlx, Nono), upstream (Sox2) and downstream (Nono). **F-I.** Upset plots identifying common and unique signatures of 7 *Evf2*-RNPs (Sox2, Smarcc2, Smarca4, Nono, Dlx, Smc1, Smc3) and 4 histone modifications (HMs) (H3K4me1/3, H3K27me3/ac) associated with *Evf2-Dlx5/6UCE* guidance, distinguish *Evf2(+)* & *Evf2(-)* regulated sites. **F.** *Evf2(+)* regulated RNPs and HMs located ±50kb to *Evf2(+)* regulated *Dlx5/6UCE*. **G**. *Evf2(+)* regulated RNPs and HMs located ±5kb to *Evf2(+)* regulated *Dlx5/6UCE.* **H.** *Evf2(-)* regulated RNPs and HMs located ±50kb to *Evf2(-)* regulated *Dlx5/6UCE*. **I**. *Evf2(-)* regulated RNPs and HMs located ±5kb to *Evf2(-)* regulated *Dlx5/6UCE*.

EGG plot comparisons of SVZ progenitors and Dlx5+ differentiated subpopulations identifies both changes in and maintenance of transcriptional targets and transcriptional outcomes, as indicated by green or red stars, which mark 9 shared target genes (Fig 3A, B). While transcriptional outcomes of 5/9 shared target genes (GM9825, Neurod6, Snrpg, Aldh1l1, Gm4604) are maintained, the majority of targets change during differentiation. For example, analysis of the furthest 4-gene cluster shows that Eno and Prmt8 are regulated in the SVZ, while Ccnd and Foxm1 are regulated in Dlx5+ populations. In addition, transcriptional reversal of Cdca3 and Rad51ap1 occurs during differentiation (SVZ to Dlx5+) (Fig 3A, B). Alignment of *Evf2* regulated RNP binding sites near SVZ and Dlx5+ transcriptional targets identifies regulation to within ±50kb in 28/29 SVZ targets, 26 within ±5kb (Fig S3). Examples of *Evf2* RNP recruitment associated with different types of EGG shifts include one-sided downstream RNP recruitment/long distance-across gene shift (Fig 3C), two-sided up- and downstream RNP recruitment/long distance-across gene shift (Fig 3D) and two-sided up- and downstream RNP recruitment/upstream shift (Fig 3E), supporting multi-modal mechanisms.

Gene clusters associated with equidistant or similar distance EGG shifts and RNP recruitment support functional links between long distance 3D co-regulation, RNP binding and transcriptional outcome. *Evf2* regulates RNPs to positively and negatively regulated Dlx5/6UCE sites (+ and - [Smarcc2, Sox2, Smarca4], - [Dlx], + [Nono], [Fig 1D and (Cajigas *et al*., 2021)], raising the possibility that RNP signatures may also distinguish EGG regulated transcriptional targets. Upset plots of RNPs and histone modifications identify shared and distinct signatures, with Sox2, Smarcc2 and Nono as the top recruited RNPs associated with *Evf2*-Dlx5/6UCE positive regulation (Fig 3F,G), and H3K4me3 associated with *Evf2*-Dlx5/6UCE negative regulation (Fig 3H, I). Therefore, differential *Evf2* RNP recruitment at *Evf2* regulated Dlx5/6UCE sites, supports that RNPs singly (as previously shown for the *Evf2*-Sox2 RNP (Cajigas *et al*., 2021)), or in specific signatures contribute to both 3D organizational and transcriptional roles.

### *Evf2* directly binds long-/super-long range EGGs on chr6 and genome-wide sites, associated with multiple RNP recruitment

We next asked whether *Evf2* direct binding contributes to EGG selectivity. We used CHIRPseq a crosslinking method to identify RNA binding sites in the genome (Chu et al., 2011). *Evf2* and Evf1, an alternatively spliced form, contain unique 5’ and common 3’ ends, and are functionally redundant in a subset of regulatory events, including RNA cloud formation, RNP recruitment, and transcriptional regulation (Cajigas *et al*., 2021; Cajigas *et al*., 2018). In order to identify Evf1/2-RNA binding sites in E13.5GEs, without the confounding contributions of 5’ or 3’ end RNA binding sites that are present in *Evf1^TS/TS^* or *Evf2^TS/TS^* single mutants (Cajigas *et al*., 2018), we utilize double mutant mice lacking both *Evf2* and Evf1 (*Evf1/2^TS/TS^*). *Evf1/2^TS/TS^* contain transcription stop insertions into Evf-exon1 and exon 3 (as described, manuscript in preparation), providing an important negative control in ChIRP experiments. CHIRPseq comparing *Evf1/2^+/+^* and *Evf1/2^TS/TS^* E13.5GEs identifies 147 *Evf1/2* RNA BSs (RBSs) across the genome (purple circles, Fig 4A). On autosomes, the highest number of RBSs are located on chr11 (22), and lowest on chr18 (3). On the X chromosome, there are 3 RBSs, whereas none are detected on the Y chromosome. Since *Evf1/2* is transcribed on chr6, it was expected that the majority of RBSs would be detected on chr6. However, only 6 RBSs are detected on chr6 at 41-129Mb distances from Dlx5/6UCE and *Evf1/2* transcription (Fig 4B), near long-range transcriptionally regulated targets (RBSs 1-3, clustered at Pdia4 (Fig 4C), RBS 4 at Creb5 (Fig 4D), and RBS 5 at Suclg2 (Fig 4E)) and super-long range target (RBS 6 at Grin2b (Fig 4F)). Analysis of *Evf2* regulated RNP binding and Dlx5/6UCE guidance associated with chr6 RBSs support a role for direct RNA binding in long distance regulatory events (Fig 4C-F). This finding is also consistent with our previous report that *Evf2* preferentially regulates long distance Dlx5/6UCE guidance across chr6, at distances >1Mb distances from the enhancer (Cajigas *et al*., 2018). As with *Evf2* regulated EGG targets, Sox2 and Smarcc2 are the top RNPs recruited to 147 RBSs, while Dlx and Nono are inhibited from binding (Upset plots, Fig 4G). HOMER de novo motif analysis identifies two motifs, including G-rich stretches that can potentially form G-quadruplexes (as shown for 4/6 chr6 RBSs (Fig 4B, H, I)). Furthermore, sequence alignments between RBSs on chr6 identifies 174 bases of 100% identity between RBS 1 and 3 (Fig 4J), supporting the possibility of RNA mediated DNA-DNA base pairing interactions (DNA looping) in EGG selectivity.

**Figure 4.**
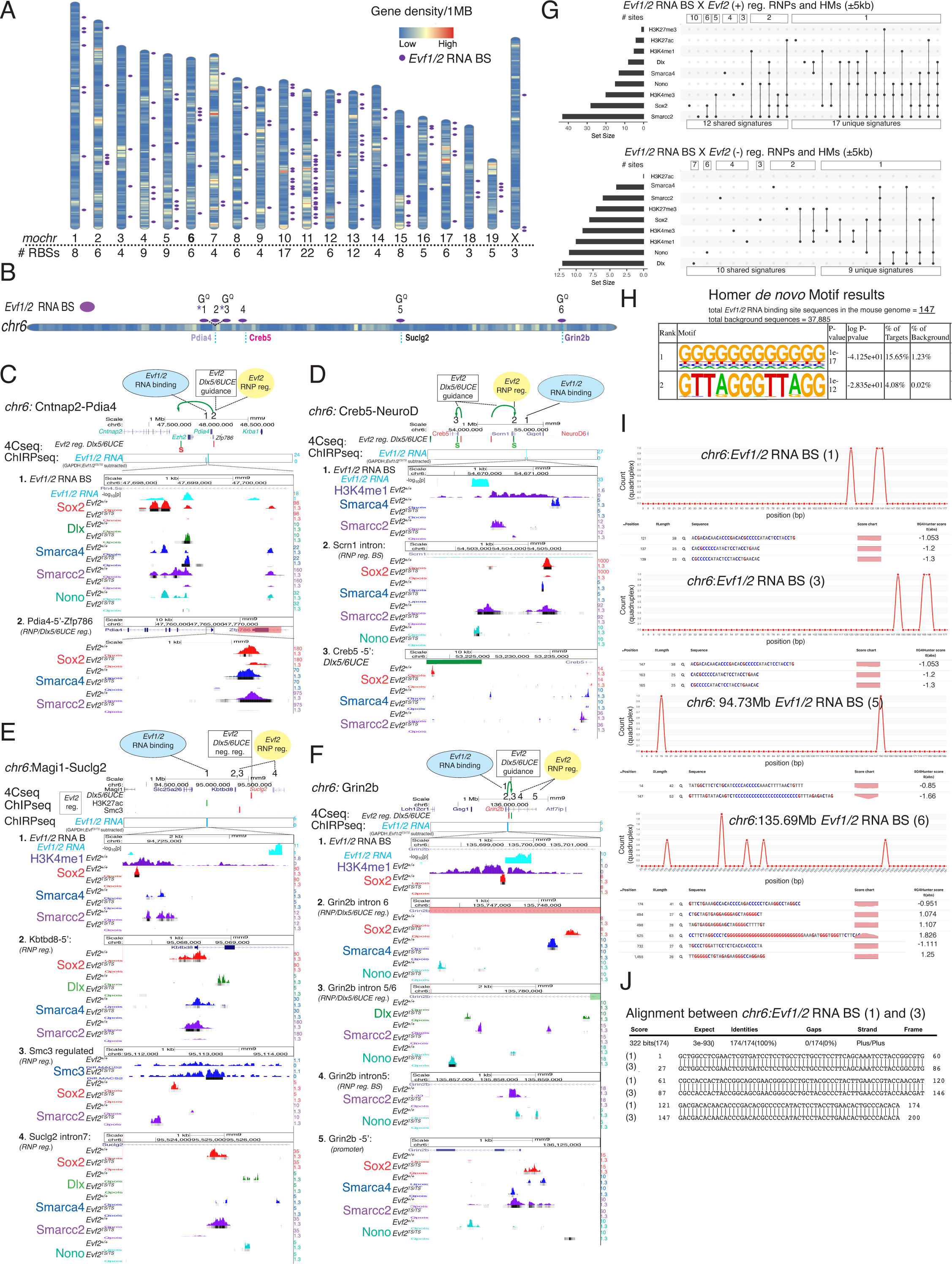
*Evf1/2* RNA direct binding sites on *chr6* localize to long- and super-long-range EGGs, multi-RNP recruitment, G-rich/quadruplex and DNA:DNA looping potential, and transcriptional regulation. **A**. ChIRPseq analysis in *Evf1/2^+/+^*E13.5 GE identifies 147 genome-wide *Evf1/2* RBSs (purple ovals) mapped onto mouse chromosomes (low to high gene density: blue to red range): from highest (22 on *chr11*), to fewest (3 on *chrX*), zero on Y (not shown). **B.** *Evf1/2* RBSs (1-6) on *chr6* map to long- and super-long range targets. **C-F**. *Evf1/2* RBS-associated *chr6* EGGs, transcriptional outcomes, RNP binding effects, and H3K4me1 marks. **C**. chr6 *Evf1/2* RBS 1-3 at (1), affects multi-RNP binding at sites (1) and at (2, Pdia4-5’), guiding *Dlx5/6UCE* from site 2 (red bar, Zfp786) to Evf2(+) reg Dlx5/6UCE (green bar) in the Cntnap2-Ezh2 intronic region (green arrow), coincident with Sox2-*Dlx5/6UCE* negative regulation (red S, from (Cajigas *et al*., 2021)); in *Evf2^TS/TS^* Cntnap2, Ezh2, Pdia4s, and Krba1 (but not Zfp786) transcripts. increase **D.** chr6 *Evf1/2* RBS 4 at (1), marked by H3K4me1, affects multi-RNP binding at site (1), and sites (2, Scrn1 intron and site 3, Creb-5’), guiding *Dlx5/6UCE* to sites 2 and 3 (green arrow, green bar), coincident with Sox2-*Dlx5/6UCE* positive regulation (green S); in *Evf2^TS/TS^* NeuroD6 and Creb5 but not Scrn1 or Ggct transcripts decrease. **E.** chr6 *Evf1/2* RBS 5 at site (1), marked by H3K4me1, *Evf2(+)* regulated H3K27ac; *Evf2* affects multi-RNP binding at site (1), and sites (2: Kbtbd8-5’, 3: *Evf2 (-)* reg Smc3, 4: Suclg2 intron7); *Evf2(-)* regulated *Dlx5/6UCE* between sites 3&4 (red line); in *Evf2^TS/TS^* Suclg2, but not Magi-Kbtbd8 transcripts decrease. **F**. chr6 *Evf1/2* RBS 6 at site (1), marked by H3K4me1; *Evf2* affects multi-RNP binding at site (1), and sites (2: Grin2b intron6 [*Evf2(-)* reg. *Dlx5/6UCE*], 3: Grin2b intron5/6 [*Evf2(+)* reg. *Dlx5/6UCE*], 4: Grin2b intron5, 5: Grin2b-5’); *Evf2* guides *Dlx5/6UCE* from site2 to site 3 (green arrow); in *Evf2^TS/TS^* Grin2b, but not adjacent transcripts decrease. **G**. Upset plots identifying common and unique signatures of 5 *Evf2*-RNPs (Sox2, Smarcc2, Smarca4, Nono, Dlx), Smc1, Smc3 (not bound) and 4 HMs (H3K4me1/3, H3K27me3/ac) within ±5kb *Ev1/2* RBSs, distinguish *Evf2(+)* and *Evf2(-)* regulation. **H.** Homer de novo analysis identifies G-rich sequences and GTTAGGGTTAGG motif enrichment at *Evf1/2* RBSs**. I.** *chr6 Evf1/2* RBSs 1, 3, 5, 6 contain G stretches with the potential to form G-quadruplex, **J.** *chr6 Evf1/2* RBSs 1-3 contain 174 bases of 100% matched sequence with the potential to form DNA:DNA loops.

We next asked whether RBS DNA identities are limited to chr6 or are characteristic of genome-wide *Evf1/2* RBSs. A search for 100% identities of greater than 25bp lengths across all 147 RBSs identifies 69 sequences with the potential for forming 77 inter-chromosomal and 8 intra-chromosomal DNA:DNA loops. The nature of inter-chr interactions range between 20 RBS pairs (potentially interacting with one other site, yellow region Fig 5A), to chr13 RBS-8, a site that matches 7 sites (red star, Fig 5A). Distances between intra-chr interactions range from 4kb-50Mb (one pair on chr1, chr6, chr10, chr12, chrX, and three pairs on chr11, Fig S4 A-E), raising the potential for short and long distance DNA looping. A network of interactions involving chr6 RBS-2 and 5 indicate the potential for inter-chr multi-hubs (>3 chr) at chr6-RBS-5 and chr13-RBS-8 through DNA looping. As observed on intra-chr6 RBS-1 & 3, *Evf2* regulates RNP binding at both inter-chr RBSs (Fig 5C, D) and intra-chr RBSs (Fig 5E, F), and H3K4me3 at chr13 RBS-8 and chr4 RBS-4. Sites of RBSs coincident with RNP hotspots (regions of multi-RNP regulation (>3 RNPs), chr4 RBS-4 (Fig 5D) and red ovals (Fig 5E) support functional roles for RNP subsets in *Evf2* RNA binding and/or transcriptional regulation. While only 3 chrX RBSs are identified, multiple RNP hotspots and transcriptionally regulated targets are identified at a distance from RBSs, suggesting long-distance effects spreading from initial binding sites (Fig 5E, Fig S4F, G). In a gene dense region of *chr11,* 3 intra-chr interactions are possible, with RBS 17-19 (5kb distance) nested within RBS 16-21 (11.6Mb distance), supporting the possibility of nested DNA looping, and RNA mediated formation of higher order DNA structures (Fig 5F).

**Figure 5.**
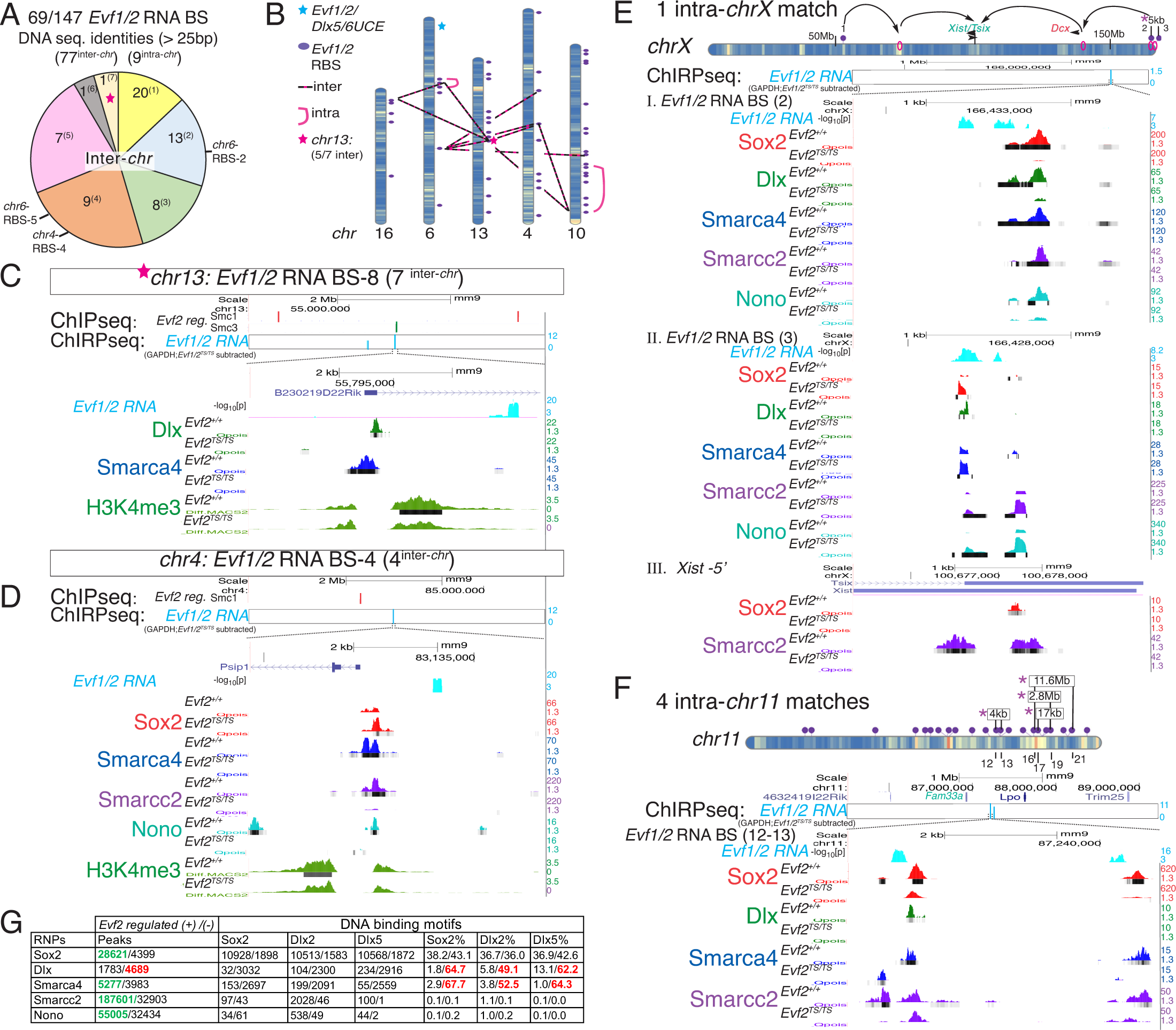
*Evf1/2* direct RNA sites identify DNA:DNA loop potential for *Evf2* intra- and inter-chromosomal regulation. **A**. *Evf1/2* RBSs (RBSs) at 100% matched DNA sequences: 69/147, >25bp, potentially forming 77 inter-chr and 8 intra-chr DNA:DNA base-pairing interactions. Pie graph showing the distribution of inter-chr interactions: most sites (20^1^) match with one other site; one site shared with 7 sites red star (1^7^). Indicated are chr6-RBS-2 (matches with 2 sites), and chr6-RBS-5 and chr4-RBS-4 (match with 4 sites). **B**. Potential inter-chr (red-black dashed) and intra-chr (pink loop), interaction network based on DNA:DNA matches at RBSs (purple ovals), *Evf1/2/Dlx5/6UCE* (blue star), chr13-RBS-8 (pink star, 5/7 inter-chr matches shown). **C.** chr13-RBS-8: *Evf2* regulated H3K4me3 and RNP binding and Smc1 (red bars) /Smc3 (green bar) regulated sites. **D**. chr4-RBS-4: *Evf2* regulated H3K4me3 and RNP binding and Smc1 (red bar) regulated site. **E.** Model for *Evf2* regulation across *chrX* (3 *Evf1/2* RBSs), intra-*chrX* match between *chrX-*RBS-2 and chrX-RBS-3 spaced 5kb apart, multi-RNP recruitment effects (pink rings, sites I (chrX-RBS-2), II (chrX-RBS-3), III (Xist-5’), transcriptional effects on *Xist, Tsix* and Dcx highlighted. **F**. Schematic of 22 RBSs across *chr11*; intra-chr11 exact matches (purple star) spacing as indicated (RBS-12-13: 4kb, RBS-16-21:11.6Mb, RBS-17-19: 17kb), identifies DNA:DNA nested potential. Cut&Run peaks identify *Evf2* regulated RNP binding in the RBS-12-13 region. **G**. *Evf2* differentially regulates RNP binding and DNA motif recognition in vivo. JASPAR analysis of Sox2, Dlx2, and Dlx5 consensus DNA binding homeobox motif enrichment at *Evf2* regulated sites, 120 fragment Cut&Run peaks in *Evf2^TS/TS^* ÷ *Evf2^+/+^*(red: decreased, green: increased). Fold change in total peaks (*Evf2* regulated binding effects), fold change in motif peaks (*Evf2* regulated motif enrichment total), fold change in % motif peaks (*Evf2* regulated motif enrichment %). Peak numbers indicated in FigS4B.

Given the associations of *Evf2* regulated RNP signatures with *Evf2* RNA binding, 3D organization on chr6, transcriptional targets, and direct effects on homeodomain RNPs Sox2 and Dlx family transcriptional activities, we next asked whether *Evf2* regulates aspects of homeodomain motif recognition in vivo. Cut&Run can distinguish between transcription factor direct (120bp fragment) and indirect (150bp fragment) binding to DNA, identifying effects on motif recognition in vivo (Meers et al., 2019b). In previous work, we showed that Sox2 consensus motifs are reduced only at Dlx and Smarca4 co-recruited sites, but not singly bound sites, supporting that *Evf2*-RNP co-recruitment affects Sox2-motif recognition, in vivo (Cajigas *et al*., 2021). Here, analysis of Sox2, Dlx2 and Dlx5 motifs in peaks from singly bound RNPs shows that *Evf2* increases Sox2 recruitment, without affecting Sox2 motif recognition (Fig 5G). However, *Evf2* decreases the number of Dlx bound sites by 2.63 fold, and decreases the % of Dlx2/5 motifs. In peaks bound by the chromatin remodeler Smarca4, an RNA/DNA binding protein with no known homeobox DNA sequence-specific binding consensus, *Evf2* decreases both Dlx2/5 and Sox2 motifs. Therefore, not only does *Evf2* affect Sox2 motif recognition at RNP co-recruited sites, but *Evf2* also inhibits Dlx recruitment by decreasing consensus Dlx2/5 DNA motif recognition, at singly-bound sites, in vivo. Of 147 *Evf1/2* RBSs, Dlx is the top negatively regulated RNP (12/147), and Sox2 the second highest positively regulated RNP (28/147), raising the possibility that *Evf2*-homeodomain motif recognition may contribute to RNA binding at a subset of sites. Taken together, these data support that *Evf2* direct binding and *Evf2*-regulated RNP binding at key sites throughout the genome may involve DNA sequence contributions including DNA:DNA base-pairing, G-quadruplex and homeobox motif recognition.

## Discussion

Chromosome capture led to the identification of 129 *Evf2* e-lncRNA regulated Dlx5/6UCE sites spanning chr6 (∼120Mb), but only 4 non-adjacent transcriptional targets across ∼40Mb, leading us to question whether *Evf2*-Dlx5/6UCE guidance is permissive, priming future transcriptional changes, or instructive, causing transcriptional changes that are masked by E13.5GE tissue heterogeneity. ScRNAseq analysis of E13.5GE subpopulations defines novel relationships between *Evf2* differentially regulated genes and multiple modes of *Evf2*-EGG regulation, supporting instructive roles. It is surprising to find that *Evf2*-GRN functional groups are both shared and unique among E13.5GE subpopulations, and that *Evf2* regulation of embryonic progenitor zone GRNs (SVZ) predicts synaptic and seizure defects detected later in adult mice lacking *Evf2* (Bond *et al*., 2009; Cajigas *et al*., 2018). Focus on *Evf2*-EGGs within 5kb of GBs (EGG-GBs), the most likely direct instructive events, reveals unexpected enhancer-gene 1D/3D relationships and subpopulation-specific effects. *Evf2* divides chr6 into activated short-range and repressed long- and super long-range EGG-GB targets, where *Evf2* regulates RNP binding at nearly every target. Sox2, an *Evf2* RNP functionally linked to *Evf2* enhancer guidance, co-regulates Dlx5/6UCE at 9/16 EGG-GBs, while *Evf2* regulation of Sox2 RNP PP formation and target gene localization is heterogenous (Cajigas *et al*., 2021). Sox2 and Smarcc2 are the top two co-recruited RNPs associated with EGG-GBs, at long-& super-long range targets (Sox2 at 7/9 (77%) and Smarcc2 at 8/9 (89%)), compared to short range targets (Sox2 at 3/8 (38%) and Smarcc2 at 6/9 (68%). Therefore, gene distances from *Dlx5/6UCE/Evf2* along chr6 level link Sox2 recruitment and transcriptional repression, consistent with our previous report of multi-level contributions of *Evf2*-Sox2 interactions (Cajigas *et al*., 2021).

While EGG-GBs are the most likely 3D events that cause transcriptional changes, targets may change in cells where 3D regulation occurs or subsequently in differentiated populations. EGG-GB transcriptional effects are identified in proliferative zones where *Evf2* is first activated, as well as in differentiated populations continuing to express *Evf2* at high levels (Lhx6), supporting immediate transcriptional effects of 3D regulation. However, EGG-GB detection in populations with decreased expression levels and low percentages of *Evf2*+ cells (Isl1/NeuroD6) (Fig 2, 6H), support long lasting or downstream effects in cells after exiting the SVZ. Given that *Evf2* is activated to the highest levels in the SVZ, the highest number of direct effects would be expected in this population. However, distribution of EGG-GBs among subpopulations does not show a preference for SVZ regulated chr6 targets (3/16). In addition, EGG clusters of 2-4 SVZ regulated chr6 target genes are distributed across chr6, raising the possibility that even genes located at ∼120Mb from the enhancer are co-regulated by Dlx5/6UCE shifts that range between 95kb-3Mb for the Eno-Cdca3-Rad51ap1-Prmt8 4-gene cluster.

While examples of enhancer RNA-gene guidance to genes within 1Mb distances are known, few reports of eRNA-EGGs span the entire chromosome, raising the question of the underlying basis for *Evf2* long-distance effects. It is possible that *Evf2* formation of Xist-like RNA clouds and multi-RNP recruitment are critical properties of chromosome-spanning EGG regulation. Also possible are critical roles of direct RNA binding sites (RBSs) at key long- and super-long-range sites, that overlap with multi-RNP recruitment, a subset characterized by G-quadruplex and DNA:DNA base pairing. We propose a model based on RBS spacing across chr6 in which RBSs facilitate specific 3D relationships within 4 chr6 regions, stabilized by the formation of region 2 (DNA:DNA base pairing between RBS1-3) (Fig 6B, i). Through stabilization of RBS defined regions, *Evf2* may facilitate enhancer hub configurations that fine-tune transcriptional outcomes of a subset of genes within a region (Fig 6B, ii accordion/nested, or 6B, iii clover). In the absence of *Evf2*, looping configurations within a region may become more relaxed, decreasing enhancer distances to a subset of regional targets, consistent with chromosome capture evidence of ectopic Dlx5/6UCE interactions (Fig 6B, iv) (Cajigas *et al*., 2018). Also consistent with an RBS regional model are multimodal enhancer-gene target relationships that regulate transcription through changes in 3D relationships, rather than simply shifting enhancers closer to genes. The model supports the idea that RNP recruitment (yellow ovals) to key sites occurs during RNA-directed enhancer shifting on chr6 (intra-chr interactions, Fig 6C), enabling transcriptional effects on both EGG-GBs and long-distance EGGs. This RBS model is consistent with the finding that RBSs play a role in regional refinement rather than establishment of large-scale chromosome spanning structures.

**Figure 6.**
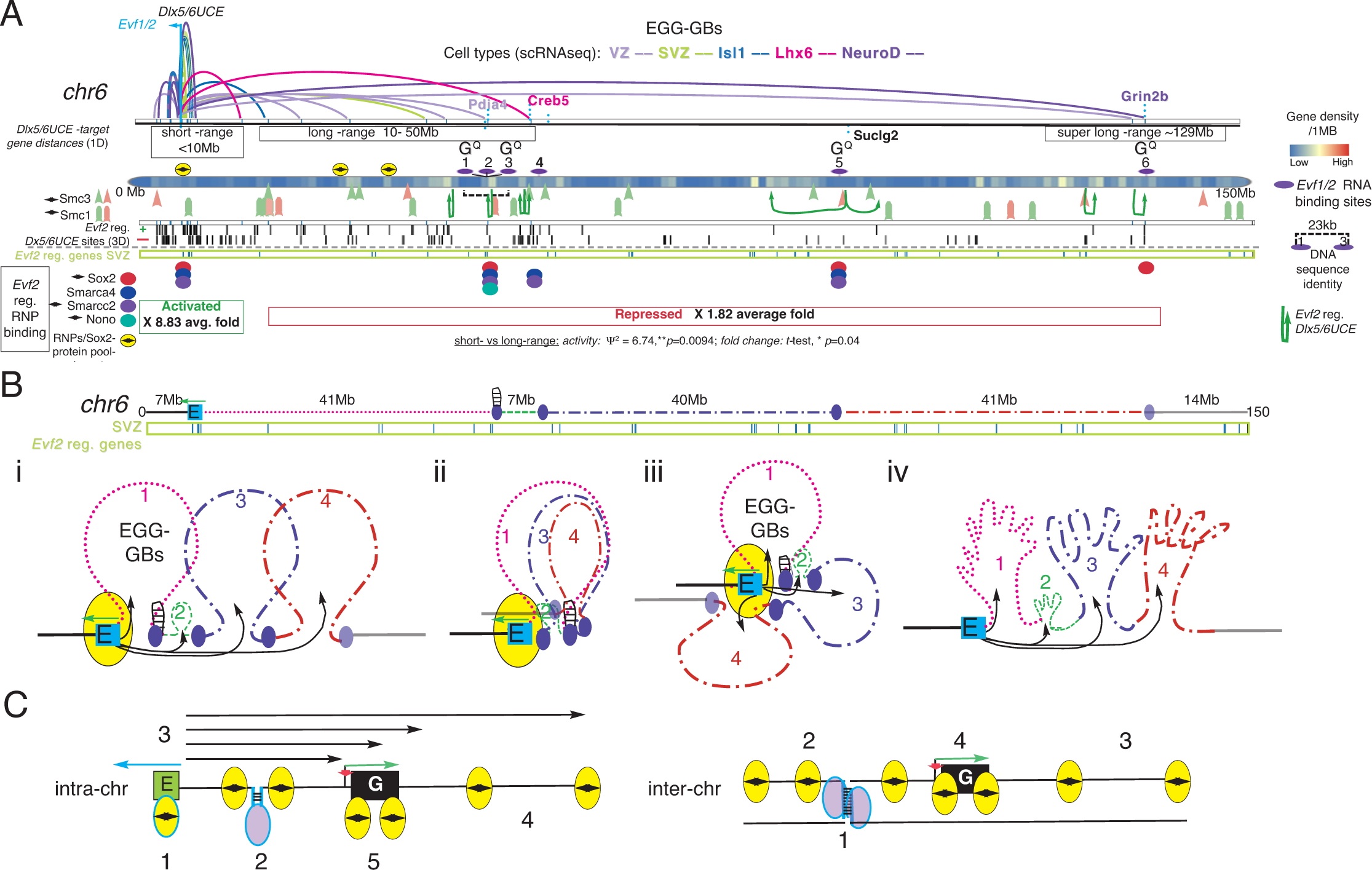
Models incorporating *Evf2* RNA direct binding, RNP recruitment, enhancer guidance and transcriptional outcome. **A-C.** Models of multi-level *Evf2* regulation on mouse chr6 resulting from RBS stabilized divisions, EGGs and RNP recruitment. **A**. Relationships between RBSs (purple ovals), DNA matches, G-quadruplex (G^Q^), RNP recruitment, EGG-GBs corresponding to scRNAseq transcriptional targets (colored loops, top), short-range activation and long-range repression domains, aligned with all EGGs (*Evf2* regulated +/- Dlx5/6UCE sites). **B**. *Evf1/2* RNA binding linear spacing and 3-D stabilized looping (1-4) of enhancer-gene groups, DNA:DNA match at RBS1-3 indicated by hairpin. i. sequential, ii. Nested/accordion loops, iii. Clover, iv. In the absence of *Evf2*, relaxed looping increases enhancer-gene targets within domains. **C**. 1. *Evf2* (blue arrow) expressed from the Dlx5/6UCE enhancer (E), recruits RNPs/PPCs (yellow oval), to E, 2. *Evf1/2* binds at chr6-RBS 1-3, facilitating DNA looping (purple oval/hairpin), recruiting RNPs, 3. Multi-modal EGG-3D changes (black arrows), 4-5. RNP recruitment and gene transcription (repression (red), activation (green). **D.** RBS stabilized DNA:DNA looping (1) in trans, affects RNP recruitment (2-4), and gene transcription (4).

### Limitations

Only 29 chr6 SVZ *Evf2*-regulated genes are identified, not only a small percentage (∼5%) of the total SVZ regulated genes, but also a small percentage of the total possible gene targets on chr6. This was surprising, indicating that *Evf2* transcriptional regulation is not enriched on chr6 over other chromosomes, and supporting the idea that *trans* mechanisms (as observed by genome wide RBSs and RNP recruitment) play a significant role in *Evf2* gene regulatory events (Fig 6D, DNA loop-predicted inter-chr interactions). Given that the extent of *Evf2* regulation of all 3D interactions (HiC) remains to be determined, we have not ruled out the possibility that cascade effects on additional enhancers, rather than Dlx5/6UCE positioning alone, contribute to transcriptional outcomes of long-distance EGGs on chr6 and in *trans*. However, previous work showed that a subset of Dlx5/6UCE-gene interactions in mouse E13.5GE (4Cseq, (Cajigas *et al*., 2018)) is conserved in human fetal brain (HiC data, (Won et al., 2016)). Furthermore, human Dlx5/6 interacts with four *Evf2*-SVZ-regulated targets (Tac1, Plxna4, Gm17829, Gm4604) and 3 Dlx5+-regulated targets (Tac1, Creb5, Gm4604) (purple H (Fig 3A, B)), supporting conservation, and therefore the significance of *Evf2*-Dlx5/6UCE guidance. Future experiments using both unbiased chromosome capture methods and genetic manipulation to distinguish between cascading and direct Dlx5/6UCE mechanisms will be important, as well as determining the role of RNA directed DNA:DNA base pairing and RNA regulated homeobox recognition to distinguish cis and trans regulatory mechanisms.

## Materials and Methods

### ScRNAseq

#### Cell Dissociation

E13.5GE dissections were collected into ice cold Leibovitz’s L-15 medium and 1% fetal bovine serum. For the single cell dissociation, papain dissociation kit, protocol II (Worthington Biochemical, LK003150). In the last step, the supernatant was removed and pelleted cells were resuspended in 1ml pre-chilled L15 medium + 1% fetal bovine serum. Cells were counted using a Luna cell counter, and viability determined to be ∼90%. The 10X Genomics microfluidics system and the 10X Genomics Chromium Single Cell 3’ Reagent Kit v3 were used for single cell separation and library preparation, following all protocol recommendations, including PCR cycles (12 cycles for cDNA amplification of 5000 cells and 14 cycles for index PCR. E13.5GE libraries from *Evf2^+/+^*and *Evf2^TS/TS^* were sequenced on the Illumina NovaSeq 6000 (Sequencing Core Facility at the La Jolla Institute).

#### ScRNAseq analysis

A custom mm9 transcriptome, in which the *Evf1* and *Evf2* transcripts were annotated as separate genes, was built using the Cellranger (version 3.1.0) mkref function. Sequencing reads were aligned to the custom transcriptome using Cellranger (version 6.1.1). A genes by cell matrix was generated from Cellranger output files within Monocle (monocle_2.24.1)(Trapnell et al., 2014) (Qiu et al., 2017a; Qiu et al., 2017b) dimension reduction was performed on the 2763 most highly variable genes determined from the transcripts/10,000 transcripts normalized matrix. Dimension reduction was performed on the top 12 principal components after regressing the effect of read depth on gene expression variance (total number of RNAs and the number of genes expressed) using uniform manifold approximation and projection (UMAP) method from the uwot package (McInnes, 2020; Melville, 2023), cells were clustered using standard approaches, with clusters assigned cell type annotations based on marker gene enrichment as performed previously (Brodie-Kommit et al., 2021; Clark et al., 2019; Lu et al., 2020). Differential expression was determined using the differentialGeneTest function within monocle, using fullModelFormulaStr = “∼genotype” and a reducedModelForulaStr = “∼genotype + Total_mRNAs” to reduce the impact of read-depth on differential expression testing. Significant differential expression was determined when q-values < 1e-5. Determination of cell cycle phase was conducted in Seurat (Seurat_5.0.1). The monocle cell dataset converted into a Seurat object, using the same variable genes as determined for input into UMAP dimension reductions. Cell cycle score and phase were determined using the mouse orthologs, as determined by biomaRt (Durinck et al., 2005; Durinck et al., 2009) of the human cell cycle genes using the CellCycleScoring function on the Seurat data object (Hao et al., 2024). Cell cycle score and phase were then transferred back to the monocle data object for subsequent analyses and plotting.

### Cut&Run

Anti-Smarcc2 (Bethyl, Cat# A301-038A; RRID:AB_817992) and anti-Nono (Bethyl, Cat#A300-587A; RRID:AB_495510) were used for Cut&Run. Cut&Run was based on methods described (Meers et al., 2019a; Meers *et al*., 2019b), and modified for E13.5GEs, as previously described (Cajigas *et al*., 2021).

#### Chromatin Isolation by RNA Purification (ChIRP)

The method described in (Chu *et al*., 2011) was followed, and modified for embryonic day 13.5 ganglionic eminences (13.5GEs). For each sample ∼16 E13.5GEs were dissected for each genotype (*Evf1/2^+/+^*, *Evf1/2^TS/TS^*) in ice cold L15, pipetted to produce a single cell suspension, pelleted for 4min, X 800 RCF, resuspended in PBS, pelleted for 4min, X 800 RCF, and resuspended in 5mL of 1% glutaraldehyde. Cells were cross-linked for 10 minutes, quenched with 500 µL of 1.25 M glycine for 5 minutes, pelleted for 5 min X 2000 RCF, 4°C, washed with PBS, pelleted for 5 min X 2000 RCF for 5min, 4°C, resuspended 1mL 4°C PBS per 8 embryos, pelleted X 2000 RCF for 3 minutes at 4°C, supernatant aspirated, and cell pellets flash frozen in liquid nitrogen and stored at -80°C.

##### Cell Lysis and Sonication

Lysis buffer (50mM Tris-HCl ph 7, 10 mM EDTA, 1% SDS +fresh .01X protease inhibitor cocktail (SIGMA), 1mM AEBSF (SIGMA A8456), and .01X RNAse inhibitor (SIGMA). Cell pellets were resuspended in 900 µL lysis buffer per 8 embryos and homogenized with a motorized pellet mixer. Cell lysate was sonicated in a Bioruptor (Diagenode) on high setting, with 24 cycles of 30 sec ON and 45 sec OFF for a total run time of 30 minutes. After sonication, 5 µL lysate was transferred to a microcentrifuge tube with 90 µL lysis buffer and 5 µL Proteinase K and incubated at 50°C for 45 minutes, and purified with ChIP DNA Clean and Concentrator Kit (Zymo Research). DNA size was checked on TapeStation, with an expected bulk of DNA around 100-500 bp. Remaining sonicated chromatin was centrifuged at 16100 RCF for 10 min, 4°C, and flash frozen in liquid nitrogen.

##### Chromatin immunoprecipitation

Chromatin was thawed at RT and 10 µL was removed for DNA INPUT. 4 mL Hybridization buffer (750 mM NaCl, 50 mM Tris-HCl pH 7, 1mM EDTA, 1% SDS, 15% formamide +fresh .01X protease inhibitor cocktail, 1mM AEBSF, and .01X RNAse inhibitor) for each pair of ChIRP reactions. 890 µL chromatin was split evenly into 3 tubes for odd, even, and GAPDH probes, and 2X volume Hybridization buffer was added. ChIRP probes were thawed at RT and 1 µL of 100µM odd, even, or GAPDH probe pool per 890 µL volume chromatin was added to separate aliquots. Aliquots were incubated at 37°C for 4 hours with end-to-end rotation. 100 µL Ampure XP beads per 100 pmol probe were prepared by washing 3 times with lysis buffer, resuspended in complete lysis buffer, added to chromatin, and incubated for an addition 30 minutes with rotation. Pelleted beads were washed 5 times with 1ml of 2X SSC, .5% SDS (+fresh 1mM AEBSF) at 37°C, incubating 5 min between each wash. Wash buffer was removed completely from the beads with the last wash.

##### DNA Isolation, libraries and sequencing

1 mL complete DNA Elution buffer (CDE: 50mM NaHCO3, 1% SDS + 10 µL of 10mg/mL RNase A and 10 µL of 10U/µl RNase H). DNA INPUT was diluted with 140 µL CDE, and incubated at 37°C for 30 minutes on a thermomixer at 300 rpm. Beads were resuspended in 150 µL of CDE, pelleted on a magnetic stand, supernatant was kept (elution 1), beads resuspended in 150 µL CDE, incubated at 37°C for 30 minutes at 300 rpm on a thermomixer, pelleted and supernatant was kept (elution 2). Elution1 and 2 were combined, 15 µL Proteinase K added, incubated at 50°C for 45 minutes at 300 rpm (thermomixer). Samples were purified with ChiP DNA Clean and concentrator kit (Zymo Research). Libraries were created with NEBNext Multiplex Oligos for Illumina (Dual Index Primers Set 1), and 10M reads from PE100 sequenced on NOVAseq. Raw sequencing reads for ChIRP datasets were aligned using bowtie2 (Langmead and Salzberg, 2012) mapper with the following settings ’--local --very-sensitive-local --no-mixed --no-discordant --phred33 -I 10 -X 700’. The datasets were mapped against the mm9 version of the mouse genome. After mapping the reads, the peak calling was performed using the MACS2 program (Zhang et al., 2008). An FDR cutoff of 0.05 was used to call the final set of peaks. The UCSC browser was used to visualize peaks, and perform alignments as shown.

##### Probe Design

Anti-sense oligo probes for *Evf2* were designed using the Biosearch Technologies’ Stellaris FISH Probe Designer. 20 total 20-mer *Evf2* probes and 10 total 20-mer GAPDH probes with a GC% ∼45% and relatively even spacing between probes were chosen and ordered with 3’-Biotin-TEG modification and HPLC purification. Probes were reconstituted at 100mM. *Evf2* probes were numbered by position from 5’ to 3’ and odd and even probe pools were created using 50 µL of each reconstituted odd and even probe respectively. 50µL of each of the 10 GAPDH probes were combined to create the GAPDH probe pool.

##### Probe Sequences for Dlx6os1 [*Evf2*]

1. CAG TGC CAT CCA ATT TGA AG
2. CTG TGA AAC TTT GGG TTC GT
3. CAG TCA GTC TTC AGA ATG GT
4. AGT CTT CTT GAA GTT GGT GT
5. TGG TTC ATC TCT GAT CTG AC
6. GTT AGC ACT CTA AGA GGT CA
7. CTA GTT CTG TGT TCT GTG AT
8. CTA CAG GGT ACA CTC AAG GA
9. GGC TTA GAG AAC ATA GCC AT
10. TAC AGT CGC ATA GCT CTT TA
11. TTG ATA AAA GAG ACC CTC CC
12. TAC CTG ATG CAT ACT GCA TA
13. ATA TTG TAT GTC AGT GCT CC
14. GAG ATA GTT AGA GCC CTT AG
15. ATA TGG ATT TGC TGA CTC CA
16. GAA CAC ACC AGA CCA TTC AT
17. CCA CCA AGA GAG TAC ATT CA
18. GAT TTC TCT TGA GGG TAC GA
19. CCT TTT TCT GTA AAC TGG CG
20. TGG TAC TCA TTT TTT CCA GG

##### Probe Sequences for GAPDH

1. AAA GGA GAT TGC TAC GCC AT
2. TTT TGA AAT GTG CAC GCA CC
3. ATT ACG GGA TGG GTC TGA AC
4. TAG AAT ACG CAT TAT GCC CG
5. ATT TAA CCT CAG ATC AGG GC
6. CAG ACC TGT GAA CTC ATT CA
7. AAT GCT TGG ATG TAC AAC CC
8. TCT CAT GTT CTT CAG AGT GG
9. ACT CAT GGC AGG GTA AGA TA
10. CCC AGT TGC TCT TAA AAG TC

#### Multiomics alignments

All scRNAseq, Cut&Run, and ChIRPseq raw and processed datasets, Metadatafile, and supplementary excels are being submitted to NCBI GEO for public access. 4Cseq chromosome capture datasets using Dlx5/6UCE as the bait were previously reported (comparing E13.5GE *Evf2^+/+^* vs *Evf2^TS/TS^* (Cajigas *et al*., 2018), and comparing Sox2fl/fl;Dlx5/6cre vs Sox2fl/fl (Cajigas *et al*., 2021)). Smc1a, Smc3, histone modification (H3K27me3, H3K27ac, H3K4me1 and H3K4me3) ChIPseq datasets from (Cajigas *et al*., 2018), and Sox2, Smarca4, Dlx Cut&Run datasets comparing E13.5GE *Evf2^+/+^*vs *Evf2^TS/TS^* were used. The UCSC Browser was used to align peaks at key regions of the mouse genome as indicated. G-quadruplex analysis was performed using https://bioinformatics.ibp.cz/#/analyse/quadruplex, based on G4hunter (Bedrat et al., 2016). Upset plots were performed using R.

## Author contributions

Conceptualization, J.D.K., B.S.C., A.C., Methodology, I.C., B.S.C, E.L., J.D.K, Software, B.S.C., A.C., S.J.K, Validation, I.C., M.L., M.B., L.C., Formal Analysis, F.S., B.S.C., A.C., S.J.K., J.D.K., Investigation, E.L.,I.C.,M.L, Data Curation, J.D.K., A.C., F.S., B.S.C., Writing-original draft, J.D.K., Writing review & editing, J.D.K., B.S.C., A.C., I.C., R.J.V., Visualization, J.D.K. S.J.K., B.S.C., Supervision, J.D.K., Project Admin. J.D.K., R.J.V., Funding acquisition, J.D.K., R.J.V.

## Acknowledgments

This work was funded by NIMH R01MH111267 and R03MH126145 to J.D.K, and RF1AG068140 to R.J.V. B.S.C is supported by an individual career development award and an unrestricted grant to the Department of Ophthalmology and Visual Sciences from Research to Prevent Blindness. Bioinformatic resources were supplemented by the National Eye Institute of the National Institutes of Health under award number P30EY002687. NovaSeq6000 was acquired through the LaJolla Institute Shared Instrumentation Grant (SIG) Program (S10OD025052). Authors have no conflicts of interest.

