## Supplemental Figures 1-4 for "Cellular heterogeneity in the developing forebrain masks transcriptional outcomes and principles of *Evf2* enhancer lncRNA-*Dlx5/6UCE*-gene guidance"

### Figure S1

A

| Sample | #Cells | #UMI | #Genes |
|---|---|---|---|
| *Evf2*$^{+/+}$(rep1) | 7587 | 5,619 ± 4,183 | 2,242 ± 1,172 |
| *Evf2*$^{+/+}$(rep2) | 4386 | 9,764 ± 5,372 | 3,217 ± 1,408 |
| *Evf2*$^{TS/TS}$ | 4321 | 9,064 ± 3,815 | 3,053 ± 884 |

B

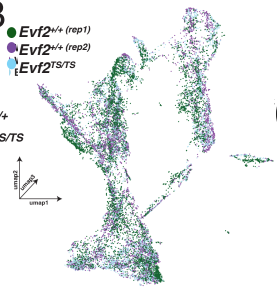


D

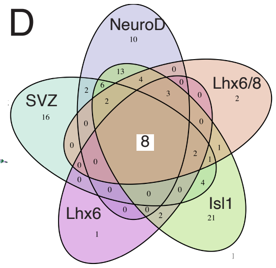


C

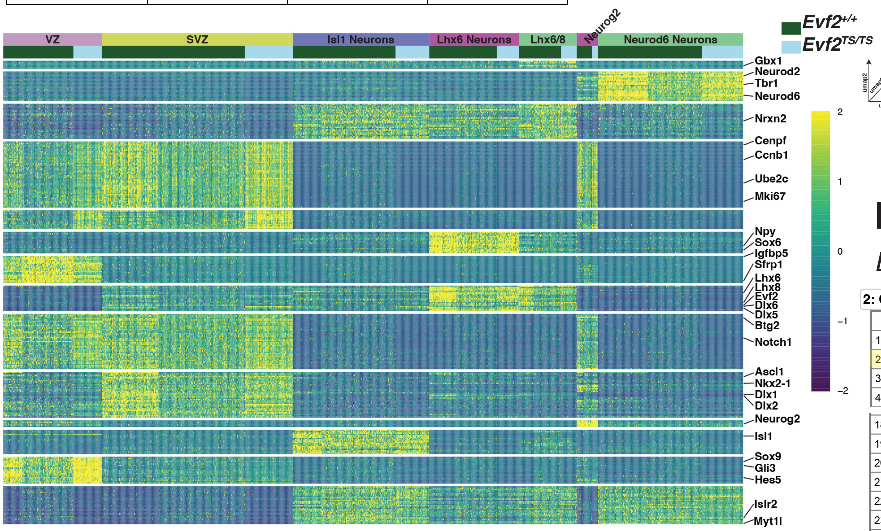


E

#### *Evf2* regulated genes (SVZ, >2-fold)

**2: GO: Biological Process** [Display Chart] 61 input genes in category / 1772 annotations before applied cutoff / 20649 genes in category

| | ID | Name | Source | pValue | FDR B&H | FDR B&Y | Bonferroni | Genes from Input | Genes in Annotation |
|---|---|---|---|---|---|---|---|---|---|
| 1 | GO:0051276 | chromosome organization | | 8.668E-8 | 1.536E-4 | 1.238E-3 | 1.536E-4 | 13 | 690 |
| 2 | GO:0034728 | nucleosome organization | | 2.660E-7 | 2.356E-4 | 1.899E-3 | 4.713E-4 | 7 | 148 |
| 3 | GO:0006334 | nucleosome assembly | | 1.525E-6 | 9.008E-4 | 7.258E-3 | 2.703E-3 | 6 | 121 |
| 4 | GO:0065004 | protein-DNA complex assembly | | 1.023E-5 | 4.530E-3 | 3.650E-2 | 1.812E-2 | 7 | 256 |

#### *Evf2* regulated genes (Lhx6, >1.5-fold)

**2: GO: Biological Process** [Display Chart] 297 input genes in category / 5168 annotations before applied cutoff / 20649 genes in category

| | ID | Name | Source | pValue | FDR B&H | FDR B&Y | Bonferroni | Genes from Input | Genes in Annotation |
|---|---|---|---|---|---|---|---|---|---|
| 1 | GO:0000278 | mitotic cell cycle | | 1.236E-58 | 6.388E-55 | 5.830E-54 | 6.388E-55 | 105 | 1089 |
| 2 | GO:1903047 | mitotic cell cycle process | | 1.399E-50 | 3.615E-47 | 3.300E-46 | 7.230E-47 | 90 | 907 |
| 3 | GO:0022402 | cell cycle process | | 1.072E-48 | 1.846E-45 | 1.685E-44 | 5.538E-45 | 110 | 1532 |
| 4 | GO:0000280 | nuclear division | | 5.289E-44 | 6.833E-41 | 6.237E-40 | 2.733E-40 | 67 | 526 |
| 5 | GO:0051301 | cell division | | 1.941E-43 | 2.006E-40 | 1.831E-39 | 1.003E-39 | 76 | 737 |
| 6 | GO:0007059 | chromosome segregation | | 9.031E-43 | 7.779E-40 | 7.100E-39 | 4.667E-39 | 64 | 488 |
| 7 | GO:0048285 | organelle fission | | 4.776E-41 | 3.526E-38 | 3.218E-37 | 2.468E-37 | 67 | 584 |
| 8 | GO:0098813 | nuclear chromosome segregation | | 1.726E-39 | 1.115E-36 | 1.018E-35 | 8.921E-36 | 55 | 372 |
| 9 | GO:0140014 | mitotic nuclear division | | 9.143E-39 | 5.250E-36 | 4.792E-35 | 4.725E-35 | 52 | 330 |
| 10 | GO:0000819 | sister chromatid segregation | | 9.896E-37 | 5.114E-34 | 4.668E-33 | 5.114E-33 | 46 | 259 |
| 11 | GO:0000070 | mitotic sister chromatid segregation | | 1.436E-36 | 6.746E-34 | 6.157E-33 | 7.420E-33 | 43 | 216 |
| 12 | GO:0051276 | chromosome organization | | 1.909E-36 | 8.223E-34 | 7.505E-33 | 9.867E-33 | 67 | 690 |

#### *Evf2* regulated genes (Lhx6/8, >1.5-fold)

**2: GO: Biological Process** [Display Chart] 82 input genes in category / 2324 annotations before applied cutoff / 20649 genes in category

| | ID | Name | Source | pValue | FDR B&H | FDR B&Y | Bonferroni | Genes from Input | Genes in Annotation |
|---|---|---|---|---|---|---|---|---|---|
| 1 | GO:0000278 | mitotic cell cycle | | 3.146E-21 | 7.311E-18 | 6.089E-17 | 7.311E-18 | 33 | 1089 |
| 2 | GO:1903047 | mitotic cell cycle process | | 3.001E-20 | 3.487E-17 | 2.904E-16 | 6.975E-17 | 30 | 907 |
| 3 | GO:0000819 | sister chromatid segregation | | 9.326E-18 | 6.247E-15 | 5.202E-14 | 2.167E-14 | 18 | 259 |
| 4 | GO:0022402 | cell cycle process | | 1.075E-17 | 6.247E-15 | 5.202E-14 | 2.499E-14 | 34 | 1532 |
| 5 | GO:0140014 | mitotic nuclear division | | 3.490E-17 | 1.622E-14 | 1.351E-13 | 8.111E-14 | 19 | 330 |
| 6 | GO:0051301 | cell division | | 4.840E-17 | 1.875E-14 | 1.561E-13 | 1.125E-13 | 25 | 737 |
| 7 | GO:0000280 | nuclear division | | 6.843E-17 | 2.272E-14 | 1.892E-13 | 1.590E-13 | 22 | 526 |
| 8 | GO:0007059 | chromosome segregation | | 2.204E-16 | 6.403E-14 | 5.333E-13 | 5.122E-13 | 21 | 488 |
| 9 | GO:0000070 | mitotic sister chromatid segregation | | 2.666E-16 | 6.884E-14 | 5.734E-13 | 6.196E-13 | 16 | 216 |
| 10 | GO:0098813 | nuclear chromosome segregation | | 3.192E-16 | 7.418E-14 | 6.178E-13 | 7.418E-13 | 19 | 372 |
| 11 | GO:0048285 | organelle fission | | 6.058E-16 | 1.280E-13 | 1.066E-12 | 1.408E-12 | 22 | 584 |
| 12 | GO:0051276 | chromosome organization | | 1.627E-15 | 3.150E-13 | 2.624E-12 | 3.781E-12 | 23 | 690 |
| 13 | GO:0007088 | regulation of mitotic nuclear division | | 3.520E-13 | 6.292E-11 | 5.240E-10 | 8.179E-10 | 12 | 141 |
| 14 | GO:0051983 | regulation of chromosome segregation | | 4.170E-13 | 6.923E-11 | 5.765E-10 | 9.692E-10 | 12 | 143 |
| 15 | GO:0051726 | regulation of cell cycle | | 1.126E-12 | 1.744E-10 | 1.452E-9 | 2.616E-9 | 27 | 1367 |
| 16 | GO:0007346 | regulation of mitotic cell cycle | | 2.871E-12 | 4.170E-10 | 3.473E-9 | 6.672E-9 | 19 | 618 |
| 17 | GO:0051783 | regulation of nuclear division | | 6.488E-12 | 8.869E-10 | 7.387E-9 | 1.508E-8 | 12 | 180 |
| 18 | GO:0007051 | spindle organization | | 1.690E-11 | 2.182E-9 | 1.817E-8 | 3.927E-8 | 13 | 248 |
| 19 | GO:0033045 | regulation of sister chromatid segregation | | 2.259E-11 | 2.763E-9 | 2.301E-8 | 5.250E-8 | 10 | 112 |
| 20 | GO:0044772 | mitotic cell cycle phase transition | | 2.523E-11 | 2.932E-9 | 2.442E-8 | 5.864E-8 | 17 | 531 |
| 21 | GO:1901992 | positive regulation of mitotic cell cycle phase transition | | 3.215E-11 | 3.558E-9 | 2.963E-8 | 7.471E-8 | 10 | 116 |
| 22 | GO:0010564 | regulation of cell cycle process | | 3.523E-11 | 3.721E-9 | 3.099E-8 | 8.187E-8 | 21 | 906 |
| 23 | GO:0051304 | chromosome separation | | 7.687E-11 | 7.767E-9 | 6.469E-8 | 1.786E-7 | 9 | 89 |
| 24 | GO:0033047 | regulation of mitotic sister chromatid segregation | | 9.436E-11 | 9.137E-9 | 7.610E-8 | 2.193E-7 | 8 | 60 |
| 25 | GO:0010965 | regulation of mitotic sister chromatid separation | | 1.612E-10 | 1.498E-8 | 1.248E-7 | 3.745E-7 | 8 | 64 |
| 26 | GO:0051306 | mitotic sister chromatid separation | | 2.352E-10 | 2.017E-8 | 1.680E-7 | 5.467E-7 | 8 | 67 |
| 27 | GO:1901989 | positive regulation of cell cycle phase transition | | 2.410E-10 | 2.017E-8 | 1.680E-7 | 5.601E-7 | 10 | 142 |
| 28 | GO:0007091 | metaphase/anaphase transition of mitotic cell cycle | | 2.431E-10 | 2.017E-8 | 1.680E-7 | 5.649E-7 | 9 | 101 |
| 29 | GO:0044784 | metaphase/anaphase transition of cell cycle | | 3.167E-10 | 2.538E-8 | 2.114E-7 | 7.361E-7 | 9 | 104 |
| 30 | GO:1901990 | regulation of mitotic cell cycle phase transition | | 6.742E-10 | 5.164E-8 | 4.301E-7 | 1.567E-6 | 14 | 406 |
| 31 | GO:0045931 | positive regulation of mitotic cell cycle | | 6.888E-10 | 5.164E-8 | 4.301E-7 | 1.601E-6 | 10 | 158 |
| 32 | GO:0033044 | regulation of chromosome organization | | 7.195E-10 | 5.225E-8 | 4.352E-7 | 1.672E-6 | 12 | 270 |
| 33 | GO:1902850 | microtubule cytoskeleton organization involved in mitosis | | 8.015E-10 | 5.645E-8 | 4.701E-7 | 1.863E-6 | 11 | 213 |
| 34 | GO:0000226 | microtubule cytoskeleton organization | | 8.419E-10 | 5.755E-8 | 4.793E-7 | 1.957E-6 | 18 | 762 |
| 35 | GO:0044770 | cell cycle phase transition | | 9.508E-10 | 6.313E-8 | 5.258E-7 | 2.210E-6 | 17 | 672 |
| 36 | GO:1905818 | regulation of chromosome separation | | 1.003E-9 | 6.474E-8 | 5.392E-7 | 2.331E-6 | 8 | 80 |
| 37 | GO:1901991 | negative regulation of mitotic cell cycle phase transition | | 1.184E-9 | 7.435E-8 | 6.192E-7 | 2.751E-6 | 11 | 221 |

F

#### *Evf2* regulated genes (SVZ, >1.5-fold)

**2: GO: Biological Process** [Display Chart] 452 input genes in category / 5635 annotations

| | ID | Name | pValue | Bonferroni | Genes from Input |
|---|---|---|---|---|---|
| 1 | GO:0031175 | neuron projection development | 1.180E-13 | 6.647E-10 | 77 |
| 2 | GO:0048666 | neuron development | 7.671E-13 | 4.323E-9 | 84 |
| 3 | GO:0050808 | synapse organization | 1.717E-12 | 9.674E-9 | 46 |
| 4 | GO:0032989 | cellular component morphogenesis | 2.951E-12 | 1.663E-8 | 65 |
| 18 | GO:0000902 | cell morphogenesis | 7.673E-10 | 4.324E-6 | 66 |
| 19 | GO:0098813 | nuclear chromosome segregation | 9.485E-10 | 5.345E-6 | 30 |
| 20 | GO:0140014 | mitotic nuclear division | 1.119E-9 | 6.304E-6 | 28 |
| 21 | GO:0061564 | axon development | 1.647E-9 | 9.280E-6 | 46 |
| 22 | GO:1902850 | microtubule cytoskeleton organization involved in mi | 1.902E-9 | 1.072E-5 | 22 |
| 23 | GO:0048667 | cell morphogenesis involved in neuron differentiatior | 2.985E-9 | 1.682E-5 | 49 |
| 24 | GO:0007059 | chromosome segregation | 3.200E-9 | 1.803E-5 | 34 |
| 25 | GO:0051276 | chromosome organization | 7.838E-9 | 4.417E-5 | 41 |
| 26 | GO:0034330 | cell junction organization | 1.096E-8 | 6.173E-5 | 50 |
| 27 | GO:0000280 | nuclear division | 2.079E-8 | 1.172E-4 | 34 |
| 28 | GO:0050890 | cognition | 2.297E-8 | 1.294E-4 | 30 |

**4: Human Phenotype** [Display Chart] 123 input genes in category / 3625 annotations

| | ID | Name | pValue | Bonferroni | Genes from Input | Genes in Annotation |
|---|---|---|---|---|---|---|
| 1 | HP:0011182 | Interictal epileptiform activity | 1.123E-6 | 4.071E-3 | 23 | 303 |
| 2 | HP:0025373 | Interictal EEG abnormality | 1.261E-6 | 4.571E-3 | 23 | 305 |
| 3 | HP:0002266 | Focal clonic seizure | 1.619E-6 | 5.868E-3 | 8 | 35 |

**5: Mouse Phenotype** [Display Chart] 366 input genes in category / 3566 annotations

| | ID | Name | pValue | Bonferroni | Genes from Input |
|---|---|---|---|---|---|
| 1 | MP:0021009 | abnormal synaptic physiology | 1.190E-8 | 4.243E-5 | 60 |
| 2 | MP:0031391 | increased locomotor activity | 3.298E-8 | 1.176E-4 | 20 |
| 3 | MP:0004046 | abnormal mitosis | 4.698E-8 | 1.675E-4 | 17 |
| 4 | MP:0001399 | hyperactivity | 5.596E-8 | 1.995E-4 | 60 |
| 5 | MP:0003635 | abnormal synaptic transmission | 5.823E-8 | 2.076E-4 | 58 |
| 6 | MP:0002206 | abnormal CNS synaptic transmission | 2.586E-7 | 9.220E-4 | 53 |
| 7 | MP:0002882 | abnormal neuron morphology | 3.532E-7 | 1.260E-3 | 73 |
| 8 | MP:0009745 | abnormal behavioral response to xenobiotic | 7.902E-7 | 2.818E-3 | 25 |
| 9 | MP:0002184 | abnormal innervation | 9.052E-7 | 3.228E-3 | 19 |
| 10 | MP:0002064 | seizures | 1.257E-6 | 4.481E-3 | 27 |

**Figure S1.** Mouse E13.5GEs from Evf2TS/TS and Evf2+/+ scSeq analysis based on 10X Genomics platform. A. Cell numbers, UMI and genes identified in scSeq samples. B. UMAP dimension reduction of all samples. C. Heatmap (yellow, highest expression, dark blue, lowest expression) of differentially expressed genes in in major cell types. D. Venn diagram of differentially expressed genes in major cell types identifying the number of overlapping and unique gene sets. E-F. Toppgene analysis of differentially regulated genes in specific cell types. Yellow highlighted terms are consistent with published evidence. GO Chromosome organization is identified as the #1 Biological process in SVZ regulated genes (>2 fold), and highly significant in differentiated cell types Lhx6 and Lhx6/8 (>1.5 fold). In the group of >1.5fold SVZ genes, the top categories include neuron development, while human and mouse phenotypes are consistent with our published evidence of increased seizure susceptibility in Evf2TS/TS mice due to reduced synaptic inhibition in prefrontal cortex (Cajigas et al 2018) and hippocampus (Bond et al 2009).

### Figure S2

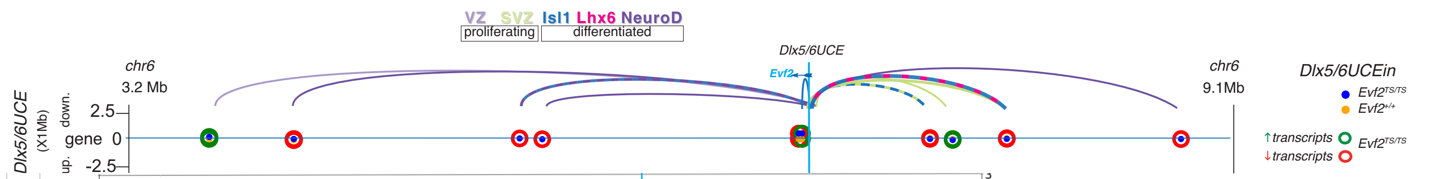


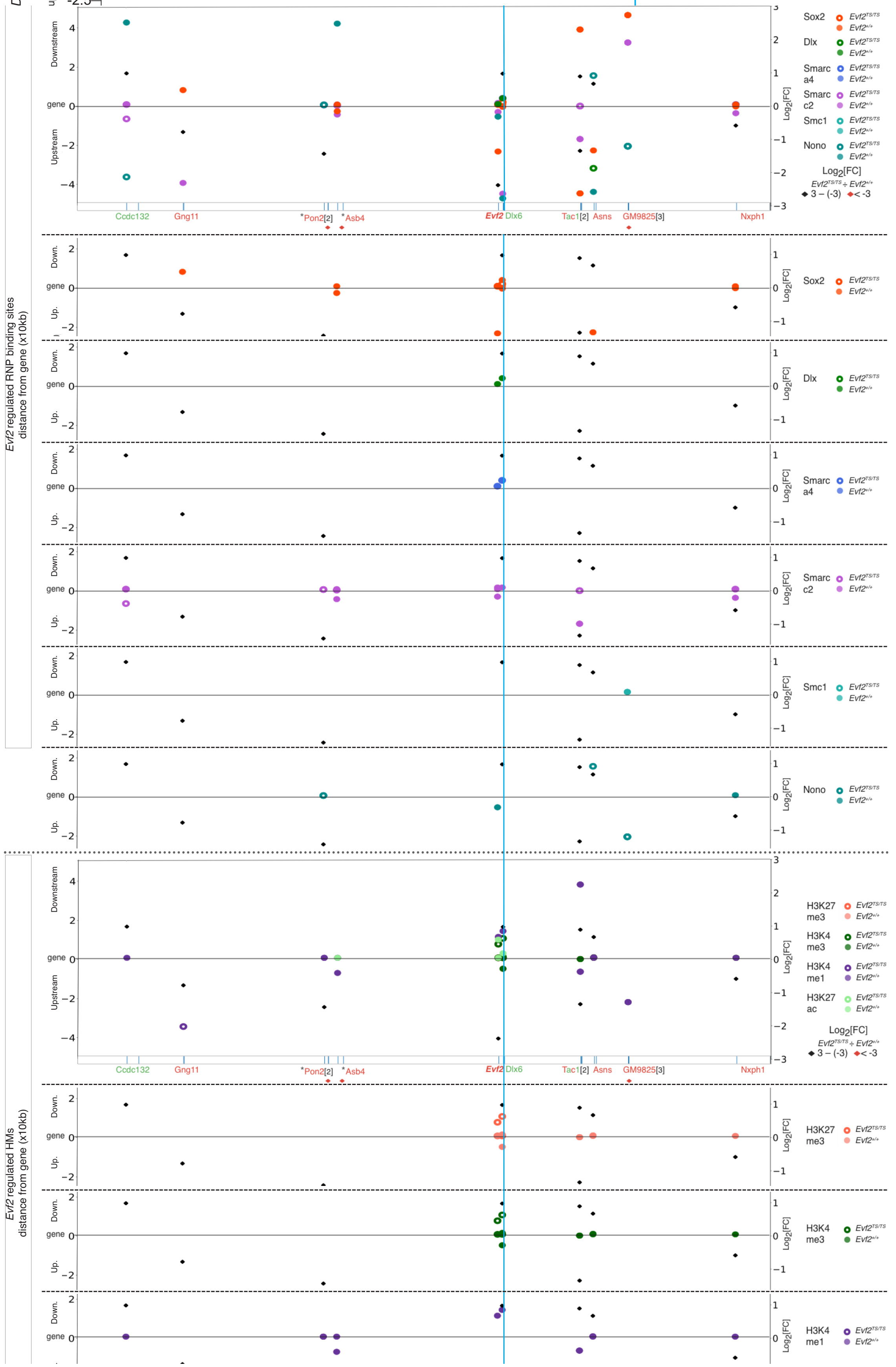


**Figure S2.** ***Evf2*-EGG-GB plots aligned with Dlx5/6UCE shifts** at each chr6 transcriptionally regulated gene target in the region between 3.2-9.1Mb, and associated Evf2 regulated Evf2-RNP binding sites are shown. Colored loops correspond to cell-type specific transcriptional regulation. Dashed loops correspond to regulation in multiple cell types (Tac1, 2 cell types, GM9825, 3 cell types). Y-axis: Evf2(+) regulated Dlx5/6UCE position (yellow dots), Evf2(-)regulated Dlx5/6UCE position (blue dots), distance from the gene (0). Transcriptional outcomes in Evf2TS/TS: shifts associated with decreased gene expression (red rings), shifts associated with increased gene expression (green rings). Right Y axis: log2 (fold change in gene expression), black diamonds (-3 - +3), red diamonds (<-3). Evf2 regulated RNP binding (Smarcc2, Smarca4, Dlx, Sox2, Nono, Smc1 (no Smc3 regulated sites): solid ovals (Evf2(+) regulated binding), rings (Evf2(-) regulated binding). Sites of Evf2 regulated histone modifications (H3K27me3, H3K4me3, H3K4me1, H3K27ac) are indicated in the bottom graph.

### Figure S3

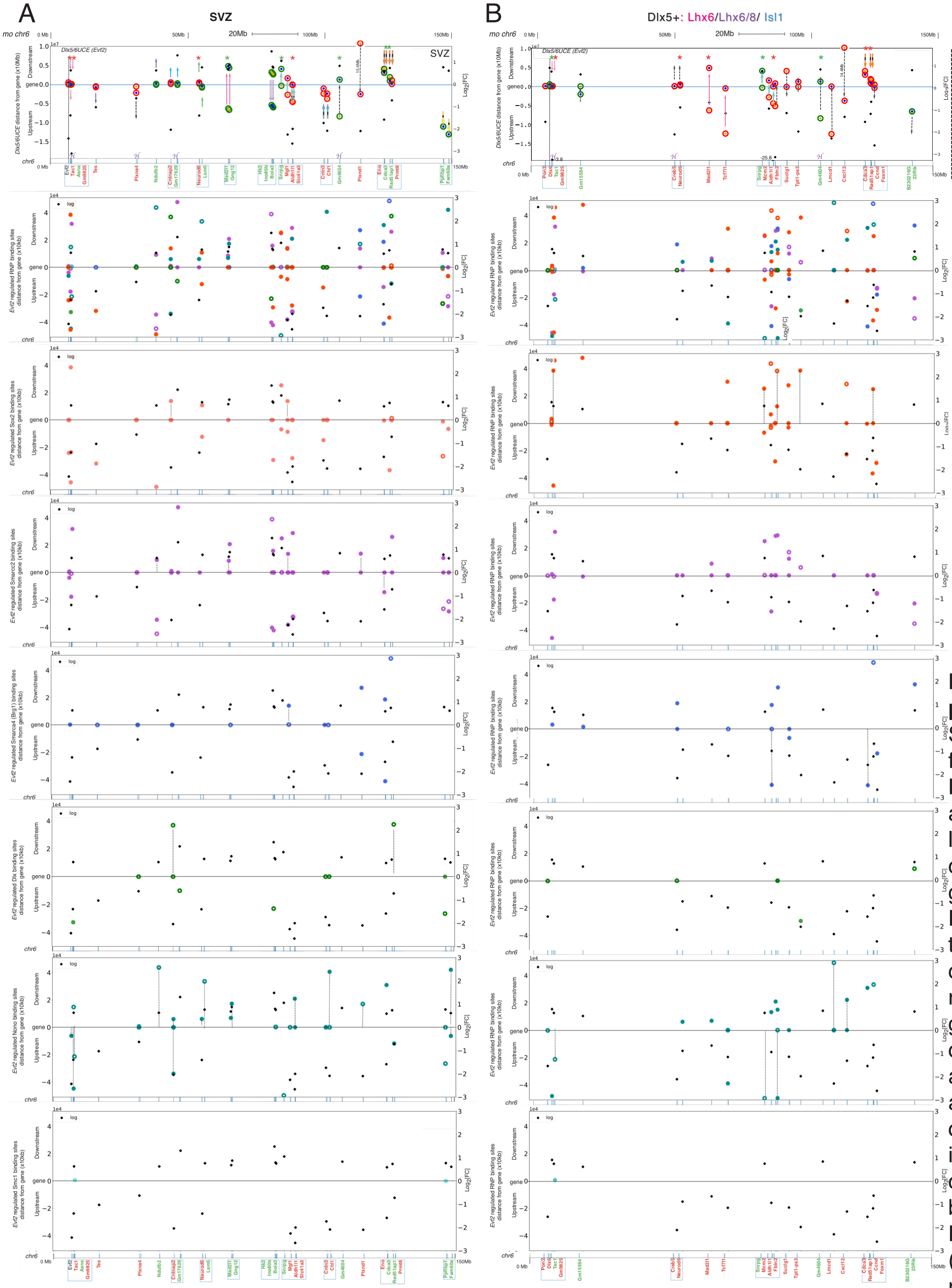


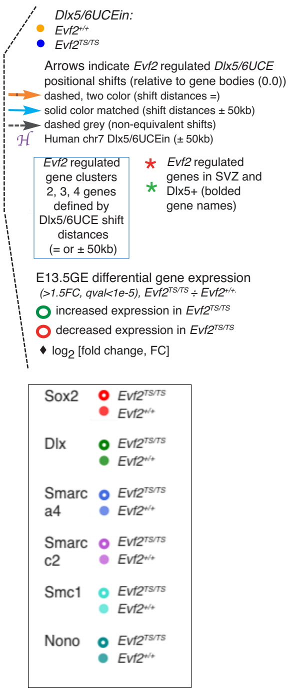


**Figure S3. Evf2-EGG plots** of left branch (A) SVZ and (B) Dlx5+ differentiated (Lhx6, Lhx6/8, Islet1) cell types at transcriptionally regulated target genes on chr6 (X-axis, 0-150Mb: gene locations), Dlx5/6UCE 3D-distances from gene bodies on Y axis (x10Mb). Arrows indicate Evf2-guided Dlx5/6UCE shift directions towards or away from gene bodies; arrow colors reflect shift distances, (as indicated in legend). Gene clusters (2-4) identified by equidistant Dlx5/6UCE shifts (dashed two color arrows) or shifts within 50kb (solid color arrows) are boxed. Conservation with human developing forebrain chr7 Dlx5/6UCE interaction sites identified by HiC (Won et al., 2016) are labelled as H. Y-axis: Evf2(+)regulated Dlx5/6UCE position (yellow dots), Evf2(-)regulated Dlx5/6UCE position (blue dots), distance from the gene (0). Transcriptional outcomes in *Evf2$^{TS/TS}$*: shifts associated with decreased gene expression (red rings), shifts associated with increased gene expression (green rings). Right Y axis: $\log_2$ (fold change in gene expression), black diamonds (-3 - +3). Stars indicate genes that are regulated in both SVZ and Dlx5+ cell types (green, increased, red, decreased). Gene clusters are boxed. *Evf2*-EGGs (±10Mb) coincident with *Evf2* regulated multi-RNP binding of Sox2, Smarcc2, Smarca4, Nono , Smc1(±50kb). RNP regulation at equidistant sites from GBs are indicated by dotted lines.

### Figure S4

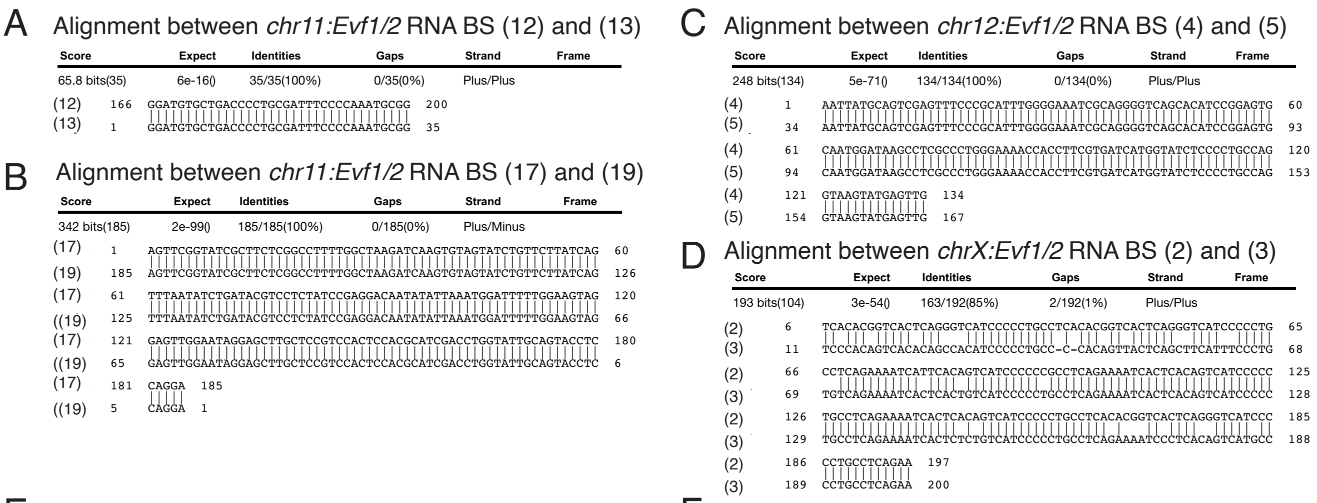


E

| *Evf1/2* RBS : Intra-chr DNA:DNA identities (>25bp) | | |
|---|---|---|
| RBS # | Location | distance |
| 2 | chr1:24338103-24338303 | 49.7Mb |
| 5 | chr1:74061897-74062097 | |
| 1 | chr6:47689894-47690094 | 23kb |
| 3 | chr6:47713254-47713454 | |
| 11 | chr10:88943028-88943228 | 32.2Mb |
| 17 | chr10:121200232-121200432 | |
| 12 | chr11:87236277-87236477 | 4kb |
| 13 | chr11:87240291-87240491 | |
| 16 | chr11:98692031-98692231 | 2.8Mb |
| 19 | chr11:101519637-101519837 | |
| 17 | chr11:101503053-101503253 | 17kb |
| 19 | chr11:101519637-101519837 | |
| 16 | chr11:98692031-98692231 | 11.6Mb |
| 21 | chr11:110302490-110302690 | |
| 4 | chr12:55839448-55839648 | 19kb |
| 5 | chr12:55820185-55820385 | |
| 2 | chrX:166427525-166427725 | 5kb |
| 3 | chrX:166432676-166432876 | |

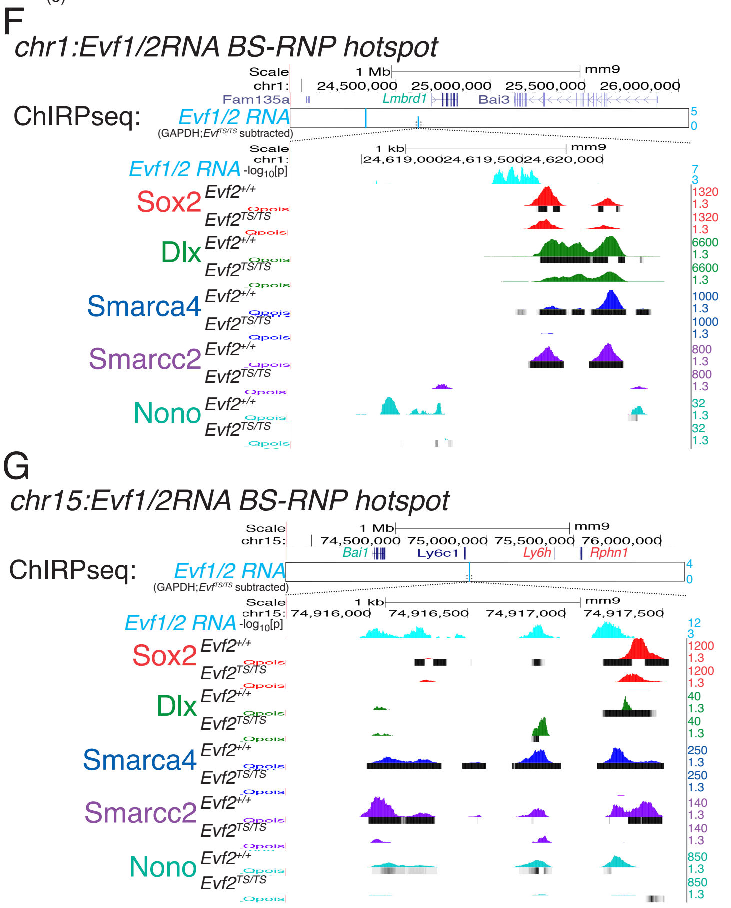


**Figure S4.** ***Evf1/2*** **RNA binding site: intra-chr DNA:DNA alignments and RNP hotspots. A-E.** 100% DNA identity (>25bp) at RBSs from ChIRPseq results. A-D. alignments of matched sites with spacing distances between 4-23kb. E, List of 9 potential intra-chr DNA:DNA matches. **F-G**. *Evf1/2* RNA binding sites overlap with multi-RNP recruitment hotspots. 120 fragment Cut&Run peaks identifying Sox2, Dlx, Smarca4, Smarcc2 and Nono binding in E13.5GE *Evf2*$^{+/+}$ vs. *Evf2*$^{TS/TS}$. Differential Qpois tracks (p<0.05) black bars below each track. UCSC Browser tracks are used.
